# Transient inhibition of type I interferon enhances CD8^+^ T cell stemness and vaccine protection

**DOI:** 10.1101/2024.06.26.600763

**Authors:** Benjamin J. Broomfield, Chin Wee Tan, Raymond Z. Qin, Brigette C. Duckworth, Carolina Alvarado, Lennard Dalit, Jinjin Chen, Liana Mackiewicz, Hiromi Muramatsu, Marc Pellegrini, Kelly L. Rogers, Woohyun J. Moon, Stephen L. Nutt, Melissa J. Davis, Norbert Pardi, Verena C. Wimmer, Joanna R. Groom

## Abstract

Developing vaccines that promote CD8^+^ T cell memory is a challenge for infectious disease and cancer immunotherapy. TCF-1^+^ stem cell-like memory T (T_SCM_) cells are important determinants of long-lived memory. Yet, the developmental requirements for T_SCM_ formation are unclear. Here, we identify the temporal window for type I interferon (IFN-I) receptor (IFNAR) blockade to drive T_SCM_ cell generation. T_SCM_ cells were transcriptionally distinct and emerged from a transitional precursor of exhausted (T_PEX_) cellular state concomitant with viral clearance. T_SCM_ differentiation correlated with T cell retention within the lymph node paracortex, due to increased CXCR3 chemokine abundance which disrupted gradient formation. These affects were due a counterintuitive increase in IFNγ, which controlled cell location. Combining IFNAR inhibition with mRNA-LNP vaccination promoted specific T_SCM_ differentiation and enhanced protection against chronic infection. These finding propose a new approach to vaccine design whereby modulation of inflammation promotes memory formation and function.

**HIGHLIGHTS:**

- Early, transient inhibition of the type I interferon (IFN) receptor (IFNAR) during acute viral infection promotes stem cell-like memory T (T_SCM_) cell differentiation without establishing chronic infection.
- T_SCM_ and precursor of exhausted (T_PEX_) cellular states are distinguished transcriptionally and by cell surface markers.
- Developmentally, T_SCM_ cell differentiation occurs via a transition from a T_PEX_ state coinciding with viral clearance.
- Transient IFNAR blockade increases IFNγ production to modulate the ligands of CXCR3 and couple T_SCM_ differentiation to cell retention within the T cell paracortex of the lymph node.
- Specific promotion of T_SCM_ cell differentiation with nucleoside-modified mRNA-LNP vaccination elicits enhanced protection against chronic viral challenge.

## INTRODUCTION

Vaccines that promote and sustain CD8^+^ T cell memory are an ongoing challenge for infectious disease and cancer immunotherapy. CD8^+^ TCF-1^+^ stem cell-like memory T (T_SCM_) cells have emerged as important determinants of long-lived T cell memory^1,2^. The induction and maintenance of T_SCM_ cells is a characteristic of vaccines regarded as exemplars of long-lived immunity^3–6^. The protective potential of T_SCM_ cells lie in their long-term persistence, high proliferative capacity and ability to generate effector CD8^+^ T (T_EFF_) cells upon rechallenge^3,6–10^. In chronic infections and cancer, a population of exhausted cells, known as precursors of exhausted (T_PEX_) cells, are marked by TCF-1 expression and are thought to be analogous to the T_SCM_ cell population identified following vaccination and acute infection^9,11–13^. T_PEX_ cells are the major cell type responding to immune checkpoint blockade^14–17^, and their numbers are predictive of patient outcome^18–20^. Given the demonstrated importance for each of these TCF-1^+^ stem-like populations for infectious disease protection and cancer immunotherapy, understanding their developmental relationship and how their generation can be specifically directed could enhance vaccine-induced protection and immunotherapy^1,21,22^.

A key determinate of CD8^+^ T cell differentiation is the site-specific environmental cues, which guide the transcriptional regulators of T cell fate^23,24^. Several studies have leveraged cytokine cues to re- invigorate existing populations of T_PEX_ cells via targeted delivery of IL-2 and CD4^+^ T cell-derived IL- 21^25–31^, or use of IL-7 and IL-15 to optimize *in vitro* differentiation of T_SCM_-like cells for chimeric antigen receptor (CAR)-T cell therapy^32,33^. In contrast, less is known about directing T_SCM_ cell differentiation for durable CD8^+^ T cell immunity. Vaccine route impacts the deposition of antigen within lymphoid tissues, which in turn modulates cell interactions and cytokines that lead to T_SCM_ cell formation^34,35^. T_SCM_ cells arise early during an inflammatory response, alongside the differentiation of effector T cells (T_EFF_)^3,36,37^. Within lymph nodes, the migration of CD8^+^ T cells into distinct niches is underwritten by transcriptional regulators. Expression of *Tbx21* (encoding T-bet) regulates CXCR3 expression and migration to the lymph node interfollicular regions (IFRs) to imprint T_EFF_ cell fate^36^. Similar to T_PEX_ cells, T_SCM_ cell differentiation is instructed by TCF-1 (encoded by *Tcf7*) which regulates CCR7 expression to retain cells in the T cell paracortex^13,36^. Thus, T_SCM_ and T_PEX_ cell differentiation occurs in distinct microenvironments to that of T_EFF_ cells^23,38^. However, how chemokine gradients are regulated to position newly activated CD8^+^ T cells is unclear.

Interferons (IFNs) are potent antiviral cytokines that belong to three major families, IFN-I, IFN-II and IFN-III^39^. The timing and location of the IFN response to viral infection is tightly coordinated^40^. Type I IFNs (IFN-Is), including IFN-α, IFN-β and IFN-ι:, all signal via the IFNα receptor (IFNAR)^39^. Dysregulation of IFN-I or their signaling pathways can exacerbate viral disease, such as influenza, measles, Herpes Simplex Virus infection, and Coronavirus Disease 2019 (COVID-19)^41–45^. Although this demonstrates IFN-Is are essential for control of viral infection, chronic induction of IFN-I counteracts positive immune responses to potentiate immune dysfunction^12,46–50^. In chronic infection, blocking IFNAR signaling directs T_PEX_ cell differentiation to reduce persistent viral load^12,48–50^. In contrast to the protective role of IFN-I during acute viral infections, expression of the only IFN-II family member, IFNγ, is a biomarker for severe disease, and in combination with TNFα can induce cell death, leading to severe tissue damage and immunopathology^40,51–53^. IFNs exert their pleotropic immune modulatory effects by inducing IFN-stimulated genes (ISGs) in multiple cell types. Although the ligands of CXCR3 – CXCL9, CXCL10 and CXCL11 – are key ISGs^54–56^, it is unknown how blocking IFNAR during viral infection impacts chemokine expression for preferential formation of T_SCM_ cells. Indeed, there is considerable overlap in the ISGs shared between distinct IFN families^39,51,57^. While the expression of CXCL9 and CXCL10 have been used as a surrogate for IFN-I expression, these ligands can be induced during acute and chronic viral infection, or recall responses by both type I and II IFNs^58–60^. Therefore, underexplored redundancy or compensation of IFN cytokines may influence chemokine expression and, in turn, CD8^+^ T cell positioning and differentiation.

As the blockade of cytokine signals that usually promote T_EFF_ cell fate, such as IL-12 and IFN-I, result in enhanced T_SCM_ cell differentiation^12,36,61^, we used temporal inhibition of IFNAR as a strategy to specifically enhance CD8^+^ stem-like T cell formation in the absence of T_EFF_ cell differentiation. We report that T_SCM_ cells transition from a T_PEX_ cellular state which coincides with viral clearance. Mechanistically, T_SCM_ cell differentiation is regulated by an unappreciated IFN-I and IFN-II interplay that augments CXCR3 chemokine expression and cell location. In combination with mRNA-lipid nanoparticle (mRNA-LNP) vaccination we demonstrate that IFNAR blockade results in enhanced vaccine efficacy, thus revealing a novel strategy to drive potent CD8^+^ T cell memory.

## RESULTS

### Early, short-term inhibition of IFN-I optimizes CD8^+^ stem-like T cell differentiation

Deficiency in IFNAR or use of IFNAR blocking antibodies increase the frequency of memory CD8^+^ T cells^12,36,39,62^. To understand the precise timing required to promote stem-like T cells, we tested how distinct IFNAR blocking schedules impacted antigen-specific P14 TCR-transgenic CD8^+^ T cell differentiation, at the peak of T cell responses 8 days (d8) following intravenous acute Lymphocytic Choriomeningitis Virus (LCMV) Armstrong infection (Figure 1A). Blocking at the time of infection and the following day (d0-1) led to the highest increase in the frequency and number of TCF- 1^+^SLAMF6^+^ stem-like T cells, that were formed in the near absence of KLRG1^+^ T_EFF_ cell differentiation (Figures 1B-1E and S1A–S1B). This was in contrast to a single dose of IFNAR blocking at the time of infection, which promoted both stem-like and T_EFF_ cell formation (Figure 1B-1E)^62^. Consistent with stem-like T cell differentiation, TCF-1^+^SLAMF6^+^ P14 cells in d0-1 IFNAR blocked mice exhibited reduced CD44, and higher CD127, CD62L, SCA-1 and programmed cell death protein 1 (PD-1) expression, similar to the T_SCM_ cells observed in control treated mice (Figure S1C).

**Figure 1.**
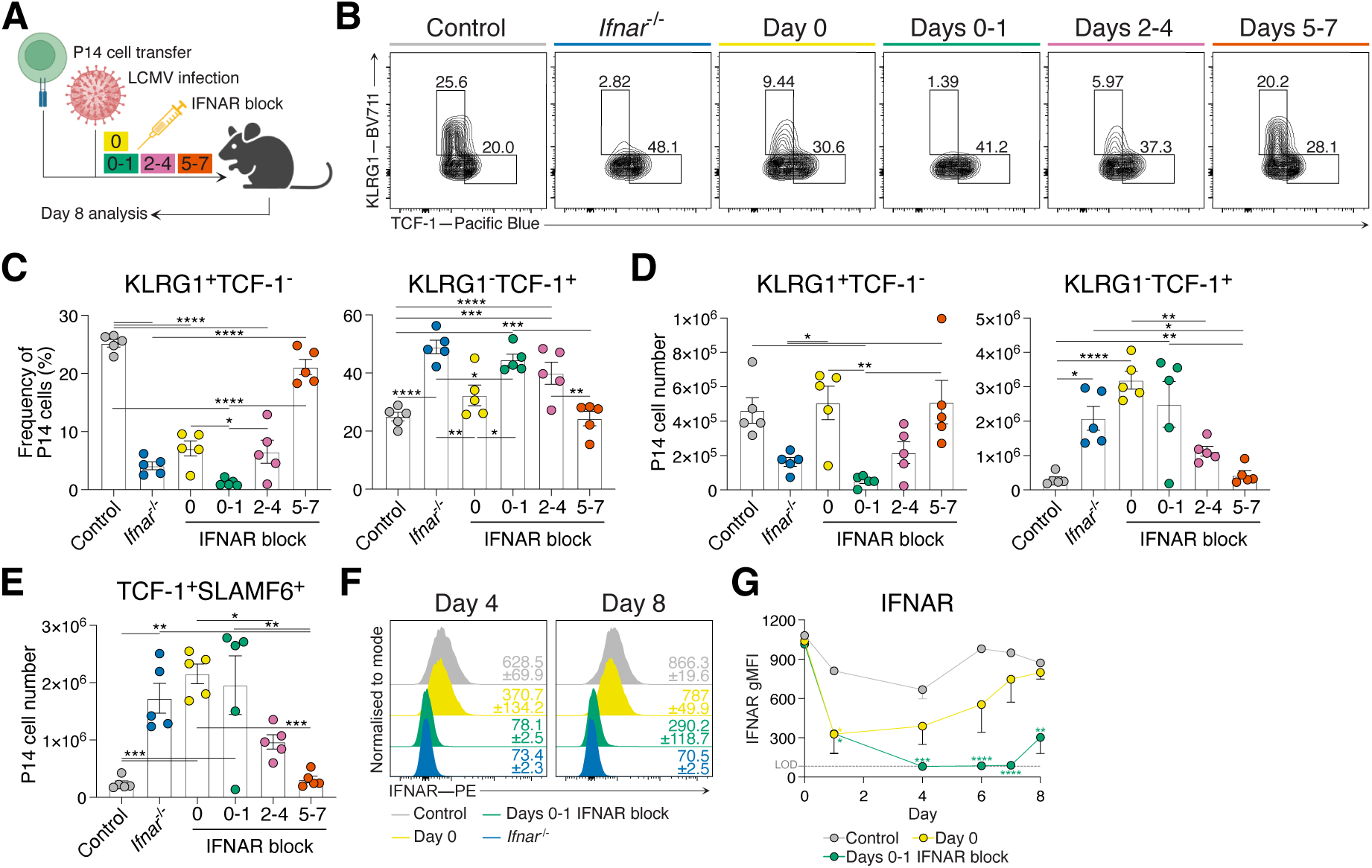
IFNAR blocking at days 0 and 1 of acute LCMV infection directs stem-like T cell differentiation. (A) Experimental scheme. P14 cells were transferred into wild type hosts prior to acute LCMV Armstrong infection and treated with indicated schedules (day 0, days 0-1, days 2-4, days 5-7) of IFNAR blocking. Peripheral lymph node P14 cells were analyzed at d8 of infection. (B-E) P14 cells generated in groups indicated in (A). Data representative of 2 independent experiments with 5 mice per group in each experiment. Each dot in (C, D, E) represent a single mouse. Data are mean ± S.E.M. Analyzed using one-way ANOVA tests. (B) Representative flow cytometry plots of P14 cells showing T_EFF_ (KLRG1^+^TCF-1^-^) and stem-like (KLRG1^-^TCF-1^+^) T cell populations. (C) Graphs summarizing frequencies in (B). (D) Graphs summarizing total P14 cell numbers of T_EFF_ and stem-like T cell subsets. (E) Graph summarizing total P14 cell number of TCF-1^+^SLAMF6^+^ stem-like T cells as shown in Supplementary Figure 1 (A). (F, G) IFNAR detection of peripheral lymphocytes following acute LCMV Armstrong infection and IFNAR blocking as indicated, or control *Ifnar^-/-^* hosts. Data representative of 3 independent experiments with 4 mice per group in each experiment. (F) Representative histograms of IFNAR expression. Average gMFI ± S.E.M. for each group is indicated. (G) Graph summarizing IFNAR gMFI. Analyzed using one way ANOVA tests. LOD dashed line indicates anti-IFNAR staining limit of detection. *p<0.05, **p<0.01, ***p<0.001, ****p<0.0001 Supplementary Figure 1 shows additional supporting data.

To understand why d0-1 IFNAR blocking led to more directed stem-like T cell formation than treatment at the time of infection alone (day 0; d0), we investigated the duration of IFNAR receptor inhibition with these treatment schedules (Figure 1F-G). Mice were infected with acute LCMV Armstrong and received IFNAR blocking on either d0, d0-1 or control treated. Throughout infection, competitive IFNAR epitope blocking was assessed by detecting binding of the same antibody clone on peripheral lymphocytes. While d0 treatment reduced anti-IFNAR binding immediately following infection, staining was regained by d4, and steadily increased to the level of the control treated mice, suggesting a single IFNAR blocking dose is insufficient to completely block the IFN-I burst at the start of acute LCMV Armstrong infection (Figure 1F-G). In contrast, in mice receiving d0-1 IFNAR block, IFNAR antibody staining remained close to the level of binding observed in *Ifnar^-/-^* cells throughout infection, reflecting complete blockade of the IFNAR epitope (Figure 1F-G). Combined, our data identify the early temporal window for IFN-I blockade during acute LCMV Armstrong infection to drive increased differentiation and total numbers of TCF-1^+^ stem-like T cells.

### Early IFNAR blockade skews stem-like T cell differentiation without establishing chronic infection

While early IFNAR blockade during acute LCMV Armstrong infection enhances the formation of stem-like T cells, IFN-I signaling is critical to overcome and clear viral infection^39,46^. To determine if d0-1 IFNAR inhibition prevented viral clearance and induced a chronic-like infection, we assessed viral load in both control treated and d0-1 IFNAR blocked acute LCMV Armstrong infection, and compared these to chronic LCMV Docile infection. At d8 of infection, acutely infected control mice had cleared LCMV Armstrong, while mice which received d0-1 IFNAR block had persistent viral load, albeit a log-fold less than mice infected with chronic LCMV Docile (Figure 2A). Unlike chronically infected mice, d0- 1 IFNAR blocked acute LCMV Armstrong infected mice completely cleared the infection by d14 (Figure 2A). The expression of the terminal exhaustion marker TIM-3 tracked with viral load (Figure 2B and S2A), and the promotion of stem-like T cells in d0-1 IFNAR blocked animals was distinct from that seen in chronically infected mice (Figures 2C and S2B). Given d0-1 IFNAR blocking during acute LCMV Armstrong infection resulted in the near absence of T_EFF_ cells, we further questioned if our observation of increased stem-like T cell differentiation was instead indicative of formation of cell exhaustion. For this, we explored the differentiation of P14 cells in control and d0-1 IFNAR blocked acute LCMV Armstrong infection and chronic LCMV Docile infection using FlowSOM dimensionality reduction analysis on an expanded number of CD8^+^ T cell differentiation markers^63^. We characterized five distinct populations from the concatenated d8 and d14 dataset using expression attributes of each FlowSOM generated population (Figures 2D and S2C-D). This revealed populations 1 and 2 to be KLRG1^+^ effector cells, distinguished by CX3CR1 expression. Population 3 had high expression of stem cell-like memory markers including SLAMF6, CD62L and CD127. Population 4 was characterized as an exhausted population, with high PD-1, TIM-3 and CXCR6 expression. To determine how stem-like T cells compared across each experimental condition, we identified SLAMF6^+^ (as a surrogate for TCF- 1 expression^64^) P14 cell populations on the tSNE map. At d8 of acute infection, SLAMF6^+^ T cells, expressed by both T_SCM_ and T_PEX_ cells, within the control treated and d0-1 IFNAR blocked mice were predominantly found in population 3 (Figure 2E). In contrast, SLAMF6^+^ T cells generated in the chronic LCMV Docile infected mice mostly overlapped with population 4 (Figure 2E), indicative of increased exhausted marker expression. By d14 following infection, almost all SLAMF6^+^ T cells within each infection setting overlaid with population 3 with the greatest number evident in the d0-1 IFNAR blocked mice (Figure S2E). Combined, these results demonstrate that stem-like T cells arising in d0-1 IFNAR blocked acute LCMV Armstrong infected mice are distinct from those in chronic infection and remain increased following viral clearance. Thus, in comparison to previous studies that demonstrate extended IFNAR blocking induces chronicity^47,48^, here we show that early, short-term IFNAR inhibition delayed viral clearance but did not establish a chronic-like infection.

**Figure 2.**
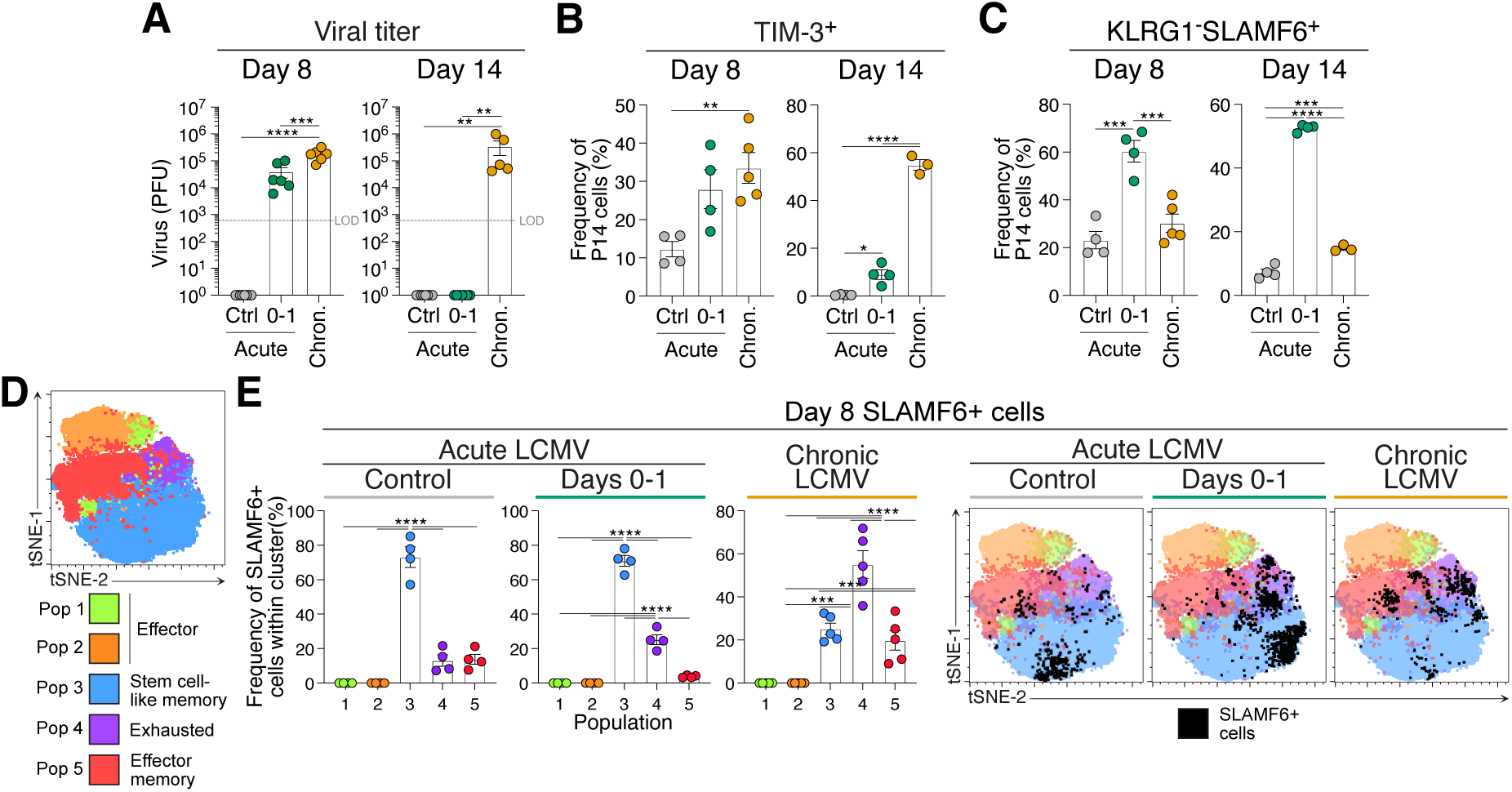
Early IFNAR blocking skews stem-like T cell differentiation without establishing chronic infection and exhaustion. Analysis of P14 cells d8 or d14 of acute LCMV Armstrong, with or without IFNAR blocking at d0-1, or chronic LCMV Docile infection. Data representative of 3 independent experiments with 4 mice per group in each experiment. Each dot in (A, B, C, E) represent a single mouse. Data are mean ± S.E.M. Analyzed using one-way ANOVA tests. (A) Graphs summarizing the plaque forming units (PFU) from viable virus in spleens of mice in each infection condition. LOD dashed line indicates viral plaque limit of detection. (B) Graphs summarizing average frequencies of TIM-3 expression on P14 cells in each indicated group. (C) Graphs summarizing frequencies of stem-like (KLRG1^-^SLAMF6^+^) T cell populations within P14 cells in each group. (D, E) FlowSOM dimensionality reduction analysis P14 cells generated in each infection condition. (D) FlowSOM dimensionality reduction analysis. tSNE plot displaying 5 FlowSOM defined cell populations and denoted identities generated from differential surface antigen expression. (E) Frequency of each FlowSOM population within d8 SLAMF6^+^ P14 cells for each infection condition, and corresponding representative overlay of SLAMF6^+^ P14 cells displayed in tSNE plots. **p<0.01, ***p<0.001, ****p<0.0001 Supplementary Figure 2 shows additional supporting data.

### T_SCM_ and T_PEX_ are transcriptionally distinct transitional cellular states concomitant with viral load

TCF-1^+^SLAMF6^+^ stem-like T cells comprise of both T_SCM_ cells that are found in acute infection and vaccination, and T_PEX_ cells, which arise during chronic infection (Figure S2E)^14,65,66^. The extended infection length observed with d0-1 IFNAR blocking during acute LCMV Armstrong infection provided a unique opportunity to investigate the developmental relationship between these cellular states. To this end, we performed paired single-cell RNAseq (scRNAseq) and CITEseq analysis, comparing d8 acute LCMV Armstrong infection (4499 cells) and chronic LCMV Docile infection (4967 cells) with d0-1 IFNAR blocked LCMV Armstrong at both d8 (5061 cells) and d14 (3102 cells). Cluster analysis revealed 15 clusters, that were broadly distinct between each infectious setting (Figure 3A-B). We then used anchor genes selected from previous studies to identify the cellular states that correspond to T_SCM_, T_PEX_ and terminally exhausted (T_EX_) cells for clusters associated with each condition (Figures 3B-C and S3A-B). Due to the close relationship between T_SCM_ and T_PEX_ cells were identified with overlapping high scores for both cellular states. Nevertheless, cluster C8, emerging from d8 acute LCMV Armstrong infection, was identified as a naturally forming T_SCM_ cell population, having a high T_SCM_ score and low T_EX_ score, in the absence of viral load (Figures 2A and 3C). Cluster C2 was the major population comprising of cells from d8 d0-1 IFNAR blocked LCMV Armstrong infection, which showed the highest score for T_SCM_ and T_PEX_ signatures. Cluster C2 also contained cells derived from chronic LCMV Docile infection, which suggested they represent T_PEX_ cells, and were distinct from clusters with high T_EX_ scores (clusters 1, 7, 12 and 3, 5, 10, 13) in each condition (Figure 3C). Despite falling in the same cluster, differentially regulated genes between these conditions indicated blocking IFNAR during priming induced genes associated with stemness, longevity and proliferation (including *Kit*, *Il7r*, *Tox*, *Bcl2*)^34,67^ (Figures 3D and S3C). Further, performing normalized enrichment score (NES) analysis with published chronic infection and cancer datasets from human and mice confirmed the identity of cluster C2 as a T_PEX_ cell state, distinct from terminal exhaustion (Figure 3E).

**Figure 3.**
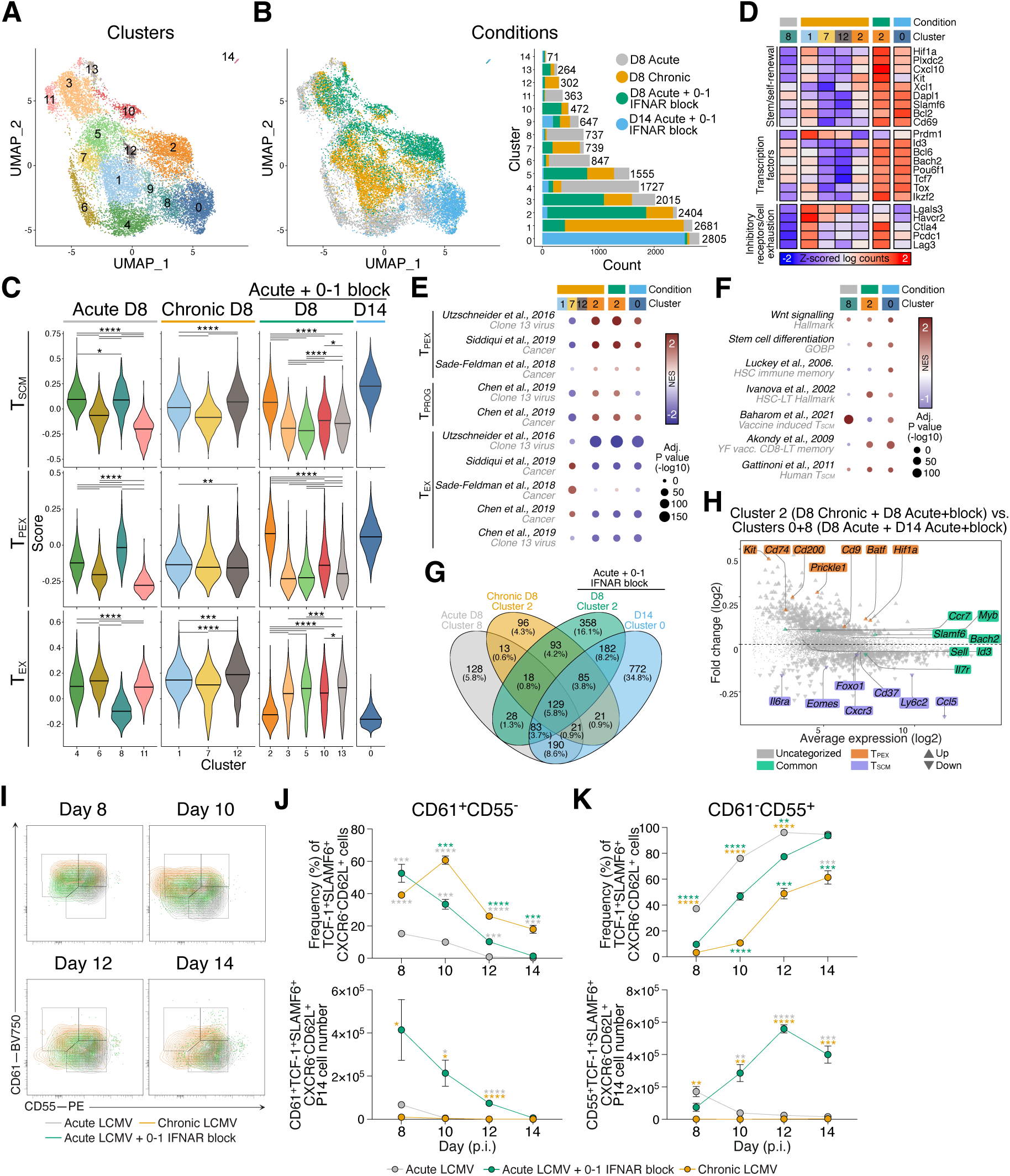
Early IFNAR inhibition promotes transitional T_PEX_ cell formation prior to establishing T_SCM_ cell population. (A-H) scRNAseq and CITEseq analysis of 17629 P14 cells from peripheral lymph nodes of d8 acute LCMV Armstrong, d8 chronic LCMV Docile, d8 and d14 IFNAR blocked acute LCMV Armstrong infection conditions. Data shows 3-4 biological replicates per condition. (A, B) UMAP projections of CD8^+^ T cells (A) based on infection condition and (C) prominence of each condition per cluster. (C) Module scores of stem cell-like memory (T_SCM_), precursor of exhausted (T_PEX_) and terminally exhausted (T_EX_)-associated genes in prominent populations from each condition. (D) Normalized mean expression heatmap of classed marker genes in selected clusters. Normalized as z-scored log counts across conditions. (E, F) Normalized enrichment score (NES) plots of enrichment of (E) T_PEX_ cell, exhausted progenitor (T_PROG_) cell and T_EX_ cell gene programs from published datasets, and (F) enrichment of hallmark Wnt signaling and stemness signatures. (G) Venn diagram reflecting distinct and intersecting gene expression in selected clusters. (H) MA plot of log fold-change versus mean expression between clusters C2 (comprising of chronic LCMV Docile and d8 IFNAR blocked acute LCMV Armstrong) and clusters C0+C8 (comprising of d14 IFNAR blocked acute LCMV Armstrong and control acute LCMV Armstrong). Marked genes represent intersections of Venn in (G), genes shared by each cluster (green), genes shared in C2 (orange), genes shared in C0+C8 (purple). Differential expression genes identified in (G) came from analysis using the *voom-limma* pipeline with duplicationCorrelation and *P* < 0.05. (I-K) P14 cell frequency and number analysis at d8 to d14 of acute LCMV Armstrong, with or without IFNAR blocking at d0-1, or chronic LCMV Docile infection. Data representative of 2 independent experiments with 3 mice per infection setting per time point. Each dot in (J, K) represent a single mouse. Data are mean ± S.E.M. Analyzed using one-way ANOVA test. **p<0.01, ***p<0.001, ****p<0.0001 Supplementary Figure 3 shows additional supporting data.

In comparison to cell states present with active infection (chronic LCMV Docile and d8 0-1 IFNAR blocked acute LCMV Armstrong), cluster C0 was the major cluster derived from d14 d0-1 IFNAR blocked acute LCMV Armstrong and had the highest T_SCM_ score of all clusters (Figures 3C and S2B). Cluster C0 expressed similar stemness genes to C2, and had decreased expression of inhibitory receptors (*Ctla4*, *Pdcd1*, *Lag3*) and increased effector molecules (*Ly6C1*, *Ccl5*), which aligned them with the naturally occurring cluster C8 T_SCM_ cells found in acute LCMV Armstrong infection (Figures 3D and S3C). NES analysis revealed that both d0-1 IFNAR blocked C0 and C2 populations exhibited enhanced Wnt signaling and stemness signatures that were predictive of stem-cell like and long-lived vaccine responses, relative to acute LCMV Armstrong memory formed without IFNAR block (Figure 3F). Analysis of intersecting genes expressed between T_PEX_ and T_SCM_ clusters, confirmed shared genes between these closely related cellular states (129 genes including *Tcf7*, *Slamf6*, *Myb*, *IL7r*, *Ccr7*) (Figure 3G-H). Pearson’s correlation analysis between T_PEX_ and T_SCM_ clusters indicated cluster C2 (chronic LCMV Docile and acute 0-1 IFNAR blocked LCMV Armstrong) were similar to each other, as were clusters C8 and C0 (comprising of cells acute LCMV Armstrong and d14 acute 0-1 IFNAR blocked LCMV Armstrong respectively) (Figure S3D). Direct comparison of these groups indicated signature genes that are either shared by and or distinguish T_PEX_ and T_SCM_ cellular states (Figure 3H). Further, we aligned CITEseq surface protein expression levels over scRNAseq UMAP clusters to indicate that CD55 and CD61 distinguish T_PEX_ and T_SCM_ populations respectively, within the TCF- 1^+^SLAMF6^+^CXCR6^-^CD62L^+^ cell population (Figure S3E-G). This allowed us to delineate the conversion from T_PEX_ to T_SCM_ cells during the course of infection. Consistent with our scRNAseq data, P14 cells in d8 IFNAR blocked acute LCMV Armstrong and chronic LCMV Docile showed a high frequency and cell number of CD61^+^ T_PEX_ cells within the TCF-1^+^SLAMF6^+^CXCR6^-^CD62L^+^ gate (Figures 3I-J and S3H-J, L). P14 cells in acute LCMV infection had a smaller frequency of these cells, which transitioned through a CD61^+^CD55^+^ population before almost entirely consisting of CD61^-^ CD55^+^ T_SCM_ cells by d12 of infection (Figures 3I-K and S3H-M). In line with delayed viral clearance, the complete transition from CD61^+^CD55^-^ T_PEX_ to CD61^-^CD55^+^ T_SCM_ was delayed to d14 in the IFNAR blocked LCMV Armstrong setting. While P14 in chronic LCMV Docile showed some transition to CD61^-^CD55^+^ T_SCM_ cells, the CD61^+^CD55^-^ T_PEX_ population was maintained at d14 post infection (Figures 3I-K and S3H-M). Combined, this analysis clarifies the distinction between T_PEX_ and T_SCM_ cellular states and identifies transcription factors and cell surface markers to unlock opportunities to track these cell fates. Further, we show that viral load critically mediates this distinction^65^. As such, early IFNAR blockade establishes a T_PEX_ cellular state that transitions into a T_SCM_ cell population with high stemness characteristics following viral clearance.

### Early IFNAR blocking increases the expression of dendritic cell derived CXCR3 chemokines

IFN-I can influence T cell differentiation, by both cell intrinsic and extrinsic mechanisms^39,46^. To untangle the direct and indirect requirement for IFNAR inhibition, we transferred wildtype or *Ifnar^-/-^*P14 cells into *Ifnar^-/-^* hosts prior to acute LCMV Armstrong infection (Figure 4A). The lack of significance between the frequency of stem-like T cells formed by wildtype and *Ifnar^-/-^* P14 cells in *Ifnar^-/-^* hosts (Figure 4B-C), indicated that IFNAR blocking primarily modulated T_SCM_ fate via altering the T cell microenvironment, rather than via T cell intrinsic mechanisms. We therefore sought to determine the underlying microenvironmental mechanisms of IFNAR blockade in driving T_SCM_ cell differentiation. We reasoned this could be due to decreased inflammation and/or via disruption of chemokine gradients that would otherwise promote T_EFF_ cell differentiation^36^. The expression of CXCR3 ligands, CXCL9 and CXCL10, are key IFN-induced environmental cues that regulate T cell positioning and function during acute and chronic viral infection, and recall responses^36,58,59,68^. We predicted that d0-1 IFNAR blocking would reduce their expression, resulting in cell retention in the lymph node paracortex to favor T_SCM_ cell differentiation^36^. To test this theory, we used the dual reporter of CXCR3 ligands, the REX3 transgenic mouse, to identify the DC chemokine cellular sources^69^ (Figure S4A). In contrast to our prediction, on d4 of acute LCMV Armstrong infection, the frequency, mean fluorescence intensity (gMFI) and total number of *Cxcl9*- and *Cxcl10*-expressing cells was increased in d0-1 IFNAR blocked REX3 mice, compared to controls (Figures 4D-E and S4B). The frequency of *Cxcl9* expression was increased in conventional type 1 and 2 (cDC1 and cDC2, respectively), while both *Cxcl9* and *Cxcl10* expression was increased in inflammatory monocyte derived DCs (moDCs) (Figure S4B). Increased chemokine expression continued to be observed in d0-1 IFNAR blocked REX3 mice at d8 of acute infection, with *Cxcl9*- and *Cxcl10*-expressing cells raised in all analyzed DC populations (Figures 4F-G and S4B). This was confirmed by light sheet fluorescent microscopy (LSFM) of whole, cleared lymph nodes d8 following acute LCMV Armstrong infection. Scaled pseudocolor images indicated increased *Cxcl9*-RFP chemokine abundance and disrupted *Cxcl10*-BFP gradient formation in d0-1 IFNAR blocked lymph nodes (Figures 4H, S4C and Supplemental Video 1).

**Figure 4.**
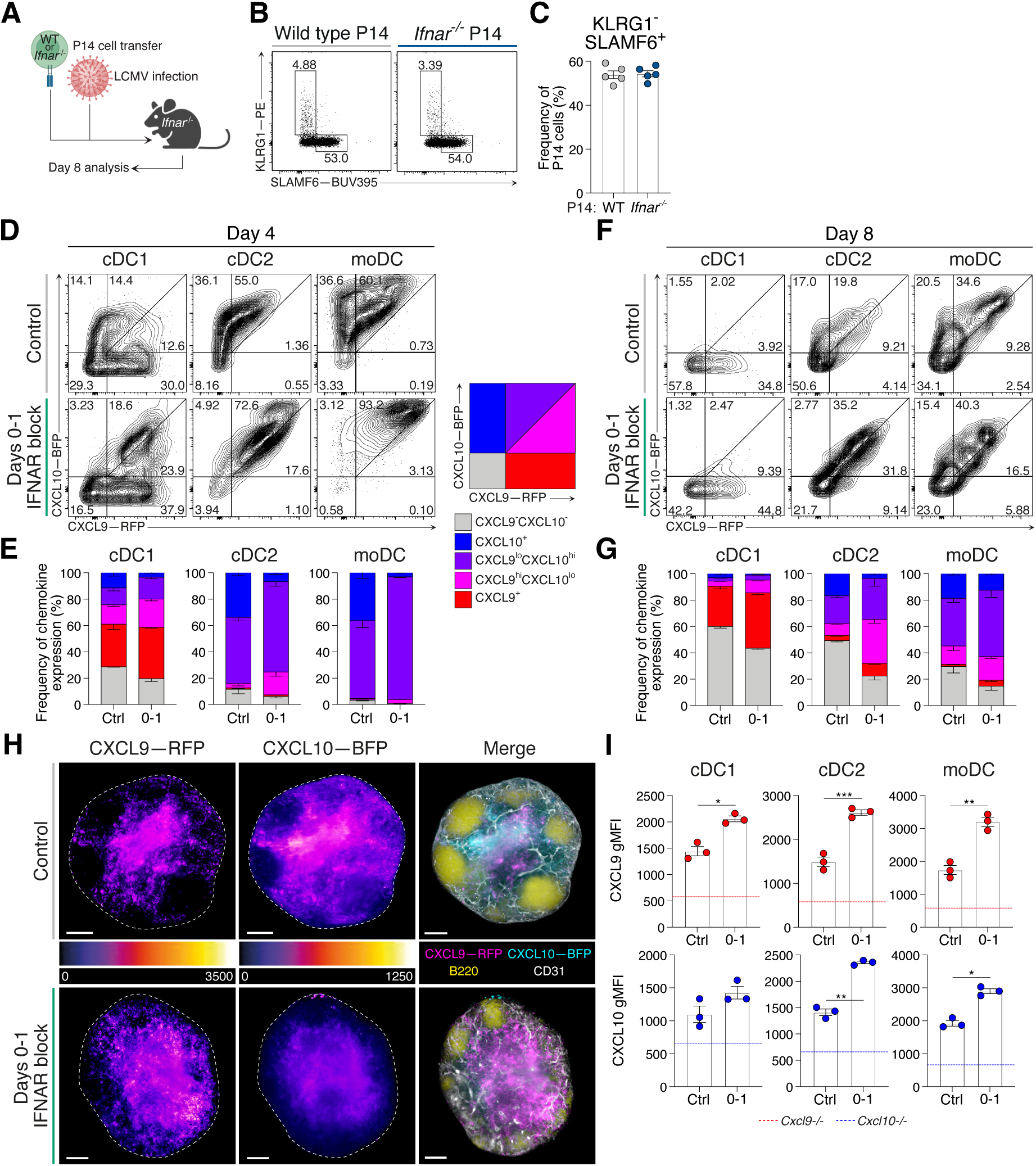
IFNAR blocking increases dendritic cell expression of CXCR3 ligands following immune challenge. (A-C) Analysis of intrinsic IFNAR signaling on the differentiation of CD8^+^ T cells. (A) Experimental scheme wildtype or *Ifnar^-/-^* P14 cell transfer into *Ifnar^-/-^* host mice prior to acute LCMV Armstrong infection and analysis at d8. (B) Representative flow cytometry plots and (C) frequency of stem-like (KLRG1^-^SLAMF6^+^) T cell populations. (D-G) REX3 reporter expression in cDC1, cDC2 and moDC populations during LCMV infection in control and d0- 1 IFNAR blocked mice. Each dot represents a single Data representative of 3 individual experiments with 3-4 mice in each group in each experiment. Data are mean ± S.E.M. (D) d4 and (F) d8 post infection representative plots showing *Cxcl9*-RFP and *Cxcl10*-BFP reporter expression in indicated DC subsets. (E, G) Graphs summarizing graded frequencies of *Cxcl9*-RFP and *Cxcl10*-BFP expression from (E, F), respectively. Graded expression summarized in key. (H) LSFM micrographs of intact REX3 lymph nodes d8 of infection in control and d0-1 IFNAR blocked conditions. Images are 200μm longitudinal slices through the lymph node center. Scale bars represent 100μm. Dashed line indicates lymph node outline. Pseudocolor FIRE LUT heat maps for each REX3 reporter (left and middle panels). Merged images (right panels) show *Cxcl9*-RFP (magenta), *Cxcl10*- BFP (cyan), B220 (B cells; yellow) and CD31 (vessels; white). Images are representative of 2 individual experiments with 4 mice in each group in each experiment. (I) Detection of CXCL9 and CXCL10 protein staining in cDC1, cDC2 and moDC populations are d4 of LCMV Armstrong infection. Graphs show average gMFI ± S.E.M. Dashed line indicates detection in DCs from *Cxcl9^-/-^* and *Cxcl10^-/-^* mice. Representative of 3 individual experiments with at least 3 mice in each group in each experiment. Each dot represents a single mouse. Statistics determined using unpaired t-tests. *p<0.05, **p<0.01, ***p<0.001 Supplementary Figure 4 shows additional supporting data.

As REX3 mice report the expression of *Cxcl9* and *Cxcl10* mRNA, it was next important to confirm that chemokine expression was increased at the protein level following d0-1 IFNAR inhibition. For this, chemokine secretion was blocked four hours prior to harvest at d4 of LCMV Armstrong infection. In line with the increased REX3 reporter expression, CXCL9 and CXCL10 showed increased gMFI following d0-1 IFNAR block, compared to control treated cDC1, cDC2 and moDC populations (Figure 4I). Therefore, d0-1 IFNAR blocking induced a paradoxical upregulation of IFN-induced chemokines following acute LCMV Armstrong infection.

Next, we sought to determine whether the increase in CXCR3 chemokine expression following d0-1 IFNAR blocking was driven by the active viral replication and high viral load (Figure 2A). REX3 transgenic mice were inoculated with either live acute LCMV Armstrong or heat-inactivated replication-deficient LCMV Armstrong. Half of each challenged cohorts were treated at d0-1 of the immune challenge and DC populations were analyzed d8 post infection. An increase in CXCR3 ligand expression, primarily *Cxcl9*-RFP, reporter expression in the d0-1 IFNAR blocked group, was observed irrespective of active or inactivated virus (Figure S4D-E). Further, these results align with decreased T_EFF_ and increased stem-like T cell formation with d0-1 IFNAR blockade with heat inactivated LCMV Armstrong infection (Figure S4F). Combined, this data indicate that d0-1 IFNAR inhibition promotes abundant CXCL9 and CXCL10 expression by cDC1, cDC2 and moDC populations, even in the absence of active viral replication, and this correlates with increased stem-like T cell formation.

### Early IFNAR inhibition leads to CD8^+^ T cell retention within the lymph node paracortex and is associated with CXCR3 desensitization

Stem-like T cell differentiation is associated with retention in the T cell paracortex in IFNAR deficient hosts^36^. This appeared at odds with the observation that d0-1 IFNAR blocking treatment increased CXCR3 chemokine expression (Figure 4). To assess how T cell location was altered with early, transient IFNAR blocking, GFP-labelled P14 cells were transferred into mice prior to acute LCMV Armstrong infection and control or d0-1 IFNAR blocking treatments. On d8 of infection, popliteal lymph nodes were cleared and analyzed by LSFM. As expected, in control treated mice, GFP-P14 cells displayed a bimodal distribution of cells, where the majority of cells were positioned surrounding the B cell follicles and deep in the inter follicular regions (IFRs), while fewer remained in the lymph node paracortex (Figure 5A and Supplemental Video 2). In d0-1 IFNAR blocked lymph nodes, the noted increase in P14 cell proliferation was evident (Figures 1C-D and 5A). Due to increased cell number, GFP-P14 cells were broadly distributed throughout the lymph node. However, compared to control treated, in d0-1 IFNAR blocked lymph nodes P14 cell location was biased towards the lymph node paracortex, consistent with previous observations in *Ifnar^-/-^* mice^36^. To quantify three-dimensional T cell location, we used eroded volume fraction (EVF) analysis, where lymph nodes are divided into 100 EVF layers of equal volume, from 0 (lymph node periphery) to 1 (lymph node center), and the density of GFP-P14 cells within each EVF was computed^36^. This method normalized for the lymphoid organ enlargement seen in d0-1 IFNAR blocked mice during acute LCMV Armstrong infection. In control treated lymph node, GFP-P14 cells located towards the lymph node periphery. In contrast, GFP-P14 cells in d0-1 IFNAR blocked lymph nodes were more homogenously distributed with a shift toward the lymph node center (Figure 5B). Using EVF analysis, we have previously identified layers of interest: 0.1-0.3, which encompasses the B cell IFRs and 0.7-0.9, within the T cell paracortex^36^. Investigating these regions in detail confirmed the location of P14 cells in d0-1 IFNAR block lymph nodes towards the center (Figure 5C). Therefore, despite increased chemokine expression, antigen specific CD8^+^ T cells in d0-1 IFNAR blocked mice are retained in the lymph node paracortex, consistent with imprinting stem-like T cell fate in this location.

**Figure 5.**
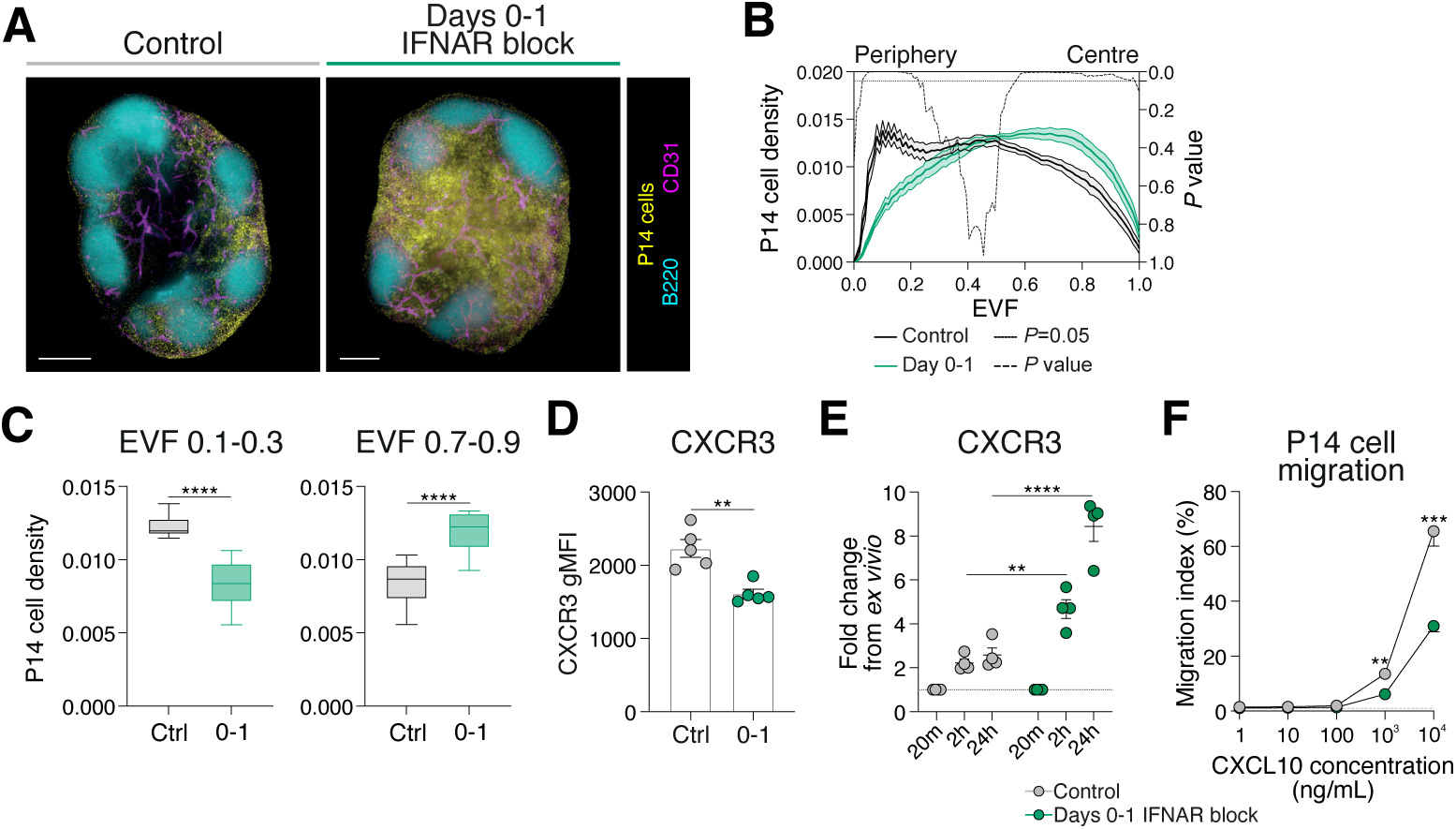
CD8^+^ T cell retention in the lymph node center is associated with CXCR3 desensitization. (A-C) LSFM micrographs of P14 cell lymph node position and density at d8 of acute LCMV Armstrong infection in control and d0-1 IFNAR blocked mice. Images and graphs representative of 3 individual experiments with 4-5 mice in each group per experiment. (A) LSFM micrographs of intact lymph nodes. Images are 200µm longitudinal slices through lymph node center. Scale bars represent 200µm. Images show GFP-P14 cells (yellow), B220 (B cells; cyan) and CD31 (vessels; magenta). (B) Graph summarizing density of P14 cells within 3D lymph node, from periphery (EVF=0) to center (EVF=1). Dashed line indicates multiple t-tests between P14 cell density in the two treatment conditions for each EVF value. (C) Graph summarizing density of P14 cells within indicated regions, interfollicular regions (EVF 0.1- 0.3) and T cell paracortex (EVF 0.7-0.9). (D-F) Analysis of P14 cells at d8 of LCMV Armstrong infection in control and d0-1 IFNAR blocked mice. Data are representative of 2 independent experiments with 4-5 mice per group in each experiment. Each dot in (D, E) represents a single mouse. Data are mean ± S.E.M. Analyzed using one-way ANOVA. (D) Graph summarizing the gMFI of P14 cell CXCR3 expression. (E) Graph summarizing the upregulation of CXCR3 on P14 cell surface at different timepoints following cell isolation. (F) P14 migratory capacity of P14 cells at different concentrations of CXCL10. **p<0.01, ***p<0.001, ****p<0.0001

The observed paracortex positioning with abundant chemokine expression with d0-1 IFNAR blockade suggested cell retention may be mediated via CXCR3 receptor desensitization. This phenomenon is well established in *in vitro* migration assays where high chemokine concentration results in G-protein uncoupling, receptor internalization and cell stasis^70^. *In vivo*, at homeostasis, desensitization of CXCR4 due to high CXCL12 sequesters lymphocytes in the bone marrow. However, how this process may control cell migration and positioning during inflammation is unknown. To investigate this, we compared the cell surface expression of CXCR3 on P14 cells in control treated and d0-1 IFNAR blocked mice infected with LCMV Armstrong. We found CXCR3 staining was decreased in mice which received d0-1 IFNAR blocking, consistent with them residing in a high chemokine concentration environment (Figure 5D). This was due, at least in part, to receptor internalization, as resting cells increased surface CXCR3 detection relative to control treated cells (Figure 5E). Consistent with this, P14 cells from d0-1 IFNAR blocked mice had reduced capacity to migrate towards CXCL10 in *ex vivo* transwell migration assays (Figure 5F). Overall, these results suggest that overexpression of chemokine following d0-1 IFNAR inhibition reduces CXCR3-directed migration to promote CD8^+^ T cell retention within the lymph node paracortex where stem-like T cell fate is imprinted.

### IFNAR blockade increases IFNγ production to tune CXCR3 chemokine expression

The upregulation of CXCR3 chemokines with IFNAR inhibition was in stark contrast to the expected regulation of these ligands^54,55^ and suggested that d0-1 IFNAR treatment may lead to an increase in an alternative IFN pathway. Consistent with this, our scRNAseq analysis indicated a counterintuitive increase in IFN signature genes (including *Cxcl10*, *Oasl1*, *Oasl2*, *Ifit3*) with IFNAR blockade (Figures 3D and S3C). NES analysis of Hallmark signatures of inflammation, IFN-I and IFN-II signaling indicated these were enhanced in IFNAR blocked samples, highlighting the considerable overlap in IFN-I and IFN-II induced signatures and unappreciated compensation between these IFN families (Figure 6A)^39,51,57^. Indeed, analysis of lymph node protein lysates showed increased expression of IFNγ in acute LCMV Armstrong infected d0-1 IFNAR blocked mice (Figure 6B), with IFNγ^+^ NK cell numbers being significantly increased (Figure S5A). Therefore, we next crossed REX3 reporter mice to generate chemokine reporters deficient in IFNAR (*Ifnar^-/-^*), IFNg (*Ifng^-/-^*) and mice double deficient in both IFNAR and IFNg (referred to as 2xIFN^-/-^ hereafter). This allowed us to investigate how deficiency in both IFN-I and IFN-II pathways impacted the expression of *Cxcl9* and *Cxcl10* following acute LCMV Armstrong infection. Similar to previous data with d0-1 IFNAR blocking (Figure 4), in *Ifnar^-/-^*lymph nodes CXCR3 chemokine expression was elevated in terms of the frequency of *Cxcl9*- RFP and *Cxcl10*-BFP expressing DCs (Figures 6C-D and S5B), gMFI of chemokine reporters and the total numbers of reporter-positive cDC1s, cDC2s and moDCs (Figure S5B). In contrast, expression of *Cxcl9*-RFP and *Cxcl10*-BFP was decreased in *Ifng^-/-^* (Figures 6C-D and S5B), with near total loss of *Cxcl9* expression. In 2xIFN^-/-^ mice, chemokine expression was largely absent. Popliteal lymph nodes from each IFN genotype were cleared and analyzed by LSFM. Again, we used scaled pseudocolor maps of individual REX3 reporter proteins to observe the relative intensity of expression in each IFN genotype. REX3 chemokine reporter expression in *Ifnar^-/-^* mice showed a striking upregulation of *Cxcl9*, with *Cxcl10* also being significantly upregulated (Figures 6E and S5C). Similar to expression profiles shown by flow cytometry analysis, *Ifng^-/-^* and 2xIFN^-/-^ mice showed a marked loss of both *Cxcl9* and *Cxcl10* (Figures 6E and S5C). In addition to DC sources, REX3 chemokines are expressed by the lymph node stromal cell compartment^36,69,71,72^. As altered REX3 expression was more pronounced between the quantified DC subsets and imaged intact lymph nodes (Figure 6C-E), we suggest a conserved IFNγ-dominant regulation of chemokine expression governs both DC and stomal cell sources. Thus, rather than the expected diminishing of inflammatory signals, early IFN-I blockade increases the expression of other inflammatory mediators, in particular IFNγ.

**Figure 6.**
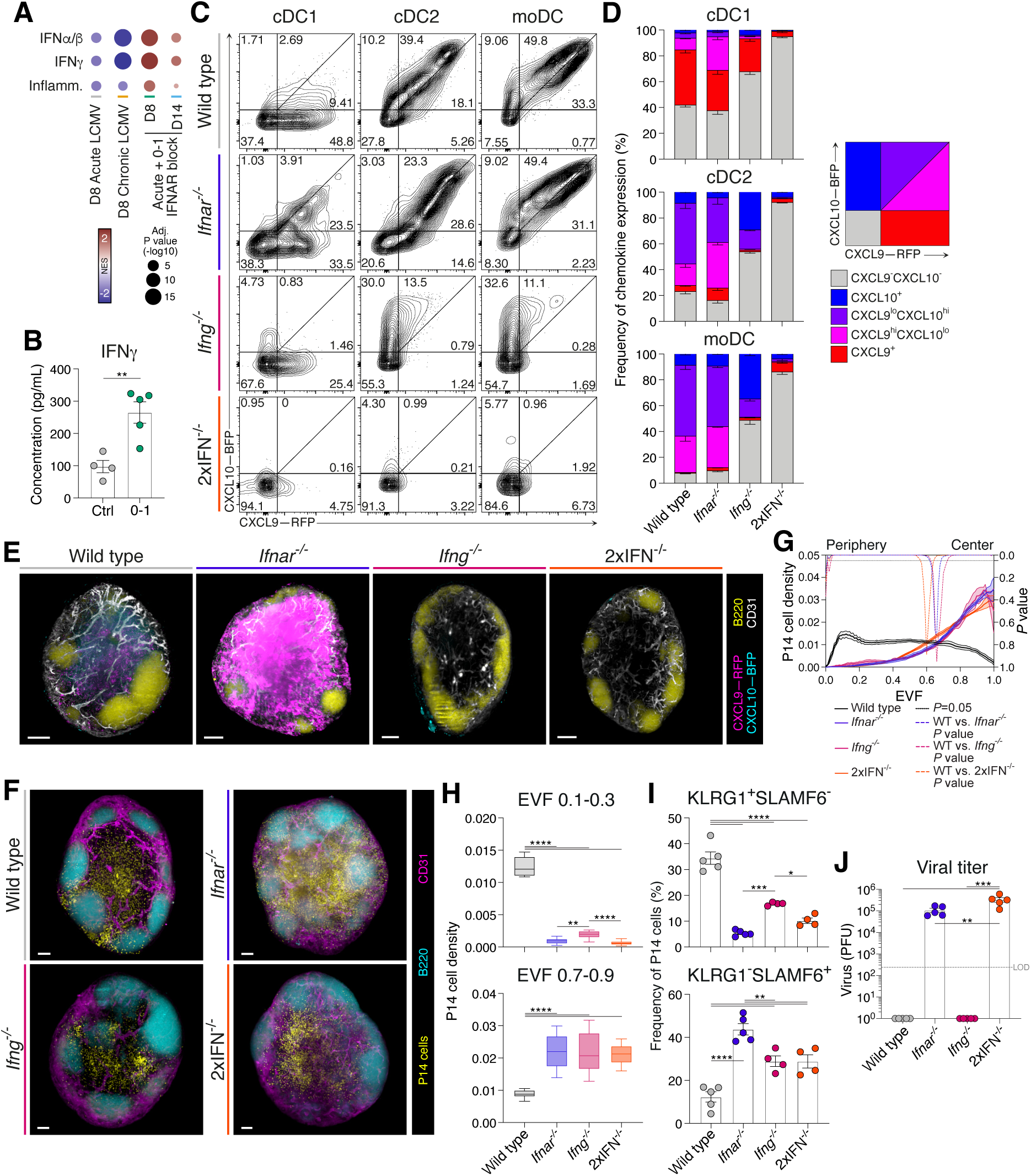
Compensatory IFNγ drives increased CXCR3 ligands in absence of IFNAR signaling. (A) NES plot of Hallmark type I and II interferon, and inflammation responses. Data generated as indicated in Figure 3. NES score represents fold change of all cells analyzed in each condition, relative to expression in all other conditions. (B) Concentration of IFNγ in lysates of lymph nodes d8 following acute LCMV Armstrong infection. Each dot represents a single mouse sample. Data are mean ± S.E.M. Statistical analysis using one-way ANOVA tests. (C, D) Expression of *Cxcl9*-RFP and *Cxcl10*-BFP in cDC1, cDC2 and moDC populations from REX3 hosts crossed to indicated IFN deficient mice at d8 of acute LCMV Armstrong infection. 2xIFN^-/-^ = *Ifnar^-/-^Ifng^-/-^*. Data representative of 3 individual repeats with at least 3 mice in each group in each experiment. (C) Representative flow cytometry plots. (D) Graphs summarizing graded frequencies of *Cxcl9*-RFP and *Cxcl10*-BFP expression from (C). Graded expression summarised in key. (E) LSFM micrographs of intact wild type, *Ifnar^-/-^*, *Ifng^-/-^* or 2xIFN^-/-^ mice REX3 lymph nodes. Images are 200µm longitudinal slices through lymph node center. Scale bars represent 100µm. Pseudocolor FIRE LUT heatmaps for each REX3 reporter (left and middle panels of each genotype). Merge images (right panels of each genotype) show *Cxcl9*-RFP (magenta), *Cxcl10*-BFP (cyan), B220 (B cells; yellow) and CD31 (vessels; white). Images are representative of 3 individual experiments with at least 3 mice in each group in each experiment. (F-H) LSFM micrographs of P14 cell lymph node position and density at d8 of acute LCMV Armstrong infection in wildtype, *Ifnar^-/-^*, *Ifng^-/-^* or 2xIFN^-/-^ mice. Images and graphs representative of 3 individual experiments with 3-5 mice in each group per experiment. (F) LSFM micrographs of intact lymph nodes. Images are 200µm longitudinal slices through lymph node center. Scale bars represent 200µm. Images show GFP-P14 cells (yellow), B220 (B cells; cyan) and CD31 (vessels; magenta). (G) Graph summarizing density of P14 cells within 3D lymph node, from periphery (EVF=0) to center (EVF=1). Dashed line indicates multiple t-tests between P14 cell density in the 3 experimental conditions against the wild type for each EVF value. (H) Graph summarizing density of P14 cells within indicated regions, interfollicular regions (EVF 0.1- 0.3) and T cell paracortex (EVF 0.7-0.9). (I) Graphs summarizing T_EFF_ (KLRG1^+^SLAMF6^-^) and T_SCM_ (KLRG1^-^SLAMF6^+^) cell populations in each of the 4 conditions. (J) Graph summarizing the plaque forming units (PFU) from viable virus in spleens of mice in each genotype. LOD dashed line indicates viral plaque limit of detection. *p<0.05, **p<0.01, ***p<0.001, ****p<0.0001 Supplementary Figure 5 shows additional supporting data.

### Stem-like T cell formation is coupled to retention within the T cell paracortex

Considering the disparity in CXCR3 chemokine expression with deficiency of IFNAR and IFNγ, we next sought to understand the impact of this on CD8^+^ T cell location. The location of GFP-labelled P14 cells was assessed in wild type and IFN deficient lymph nodes d4 following acute LCMV Armstrong infection. In control treated mice, GFP-P14 cells displayed a bimodal distribution, where the majority of cells were positioned surrounding the B cell follicles and the intrafollicular regions, while fewer remained in the lymph node paracortex (Figure 6F). Quantification of P14 cell density throughout the lymph node, within the periphery (EVF 0.1-0.3) and paracortex (EVF 0.7-0.9) again highlighted a high proportion of T cells towards the outer niches of the lymph node (Figure 6G-H). In contrast, analysis of GFP-P14 cell distribution within *Ifnar^-/-^*, *Ifng^-/-^*, and 2xIFN^-/-^ host mice lymph nodes revealed sequestering of cells within the central paracortex region (Figure 6F–H). Additionally, the increased differentiation of T_EFF_ cells in *Ifng^-/-^* mice was observed, with significantly more cells moving into the IFRs of the lymph node in this setting compared to either *Ifnar^-/-^* or 2xIFN^-/-^ hosts (Figure 6G-H). Given the dynamic regulation of CXCR3 chemokines and cell positioning, we next assessed the impact on T cell differentiation. As seen with d0-1 IFNAR blocking, IFNAR deficiency, either alone or in combination with *Ifng^-/-^* resulted in increased P14 cell proliferation at d8 of LCMV Armstrong infection (Figure S5D). Concurrent with this, IFNAR deficiency decreased T_EFF_ cells and increased stem-like T cell formation, similar to that observed with d0-1 IFNAR blocking (Figures 6I and S5E). Unlike *Ifnar^-/-^* mice, stem-like T cell formation was promoted in *Ifng^-/-^* hosts in the absence of ongoing viral load, suggesting that the inflammatory regulation of cell position plays a role in addition to increased antigen burden (Figure 6J). Combined, this data suggests that, for each IFN-deficient model, cell retention in the paracortex and increased stem-like T cell formation occurred by distinct two mechanisms. Firstly, the IFNγ-dependent increase of CXCR3 chemokines mediates gradient destruction and CXCR3 receptor internalization in *Ifnar^-/-^* (or d0-1 IFNAR blocked) lymph nodes. Secondly, the reduction or total ablation of chemokine expression in *Ifng^-/-^* and 2xIFN^-/-^ mice, limits T cell migration to the lymph node periphery (Figure S5F). In these settings, retention in the paracortex is likely mediated by TCF-1 regulated expression of CCR7^36^. Together this data underscore the importance of chemokine regulation and spatial location for the differentiation of stem-like T cells *in vivo*.

### Type I IFN inhibition promotes almost exclusive differentiation of T_SCM_ cells following mRNA- LNP vaccination

As noted, IFNAR blockade alters chemokine expression and promotes stem-like T cell differentiation independently of viral replication and viral burden (Figure S4D-F). In view of this, we reasoned that short-term IFN blockade could be leveraged in a vaccine setting for the promotion of T_SCM_ cell differentiation and enhanced immune memory. While the level of IFN-Is and IFNγ induced between viral and mRNA-LNP vaccine settings is distinct, LNPs have their own adjuvant affect and vaccine-encoded mRNA may still induce IFN-I despite nucleoside modification^73,74^. Indeed, in addition to IL-6 production, CXCL10 is also highly upregulated in lymph nodes 4 hours following LNP administration, suggesting IFNs may play a role in modulating mRNA-LNP CD8^+^ T cell responses induced by nucleoside-modified mRNA-LNP vaccines^73,74^. To test this, we generated mRNA-LNP vaccines encoding a GP33 minigene, recognized by P14 cells, to explore IFN-I and IFN-II inhibition following vaccination. Mice received adoptive cell transfers of P14 cells a day before flank thigh intramuscular vaccinations immediately prior to either or combined d0-1 IFNAR and IFNγ blocking antibodies (Figure S6A). On d8 following vaccination, mRNA-LNP-induced P14 cell differentiation was dominated by both T_SCM_ (TCF-1^+^SLAMF6^+^) and cells expressing the exhaustion marker TIM-3 (T_EX_, TIM-3^+^KLRG1^-^SLAMF6^-^) cells (Figures 7A-B and S6B-D). Distinct from our observations during acute LCMV Armstrong infection, IFNγ blockade in combination with mRNA-LNP vaccination failed to alter T_SCM_ and T_EX_ differentiation, suggesting a negligible role for IFNγ in T_SCM_ cell formation following vaccination (Figures 7A-B and S6B-C). In contrast, d0-1 IFNAR blocking alone (αIFNAR), or in combination with IFNγ blocking (α2xIFN) ablated both T_EFF_ and T_EX_ cell formation, leading to the specific promotion of T_SCM_ cells (Figures 7A-B and S6B-D). Thus, T_SCM_ cells formed ∼90% of all antigen-specific CD8^+^ P14 T cells, compared to ∼40% in control treated and IFNγ single-blocked (αIFNγ) groups. Therefore, d0-1 IFNAR inhibition in combination with mRNA-LNP vaccination drives the almost exclusive formation of T_SCM_ cells that is distinct from exhaustion. Thus, demonstrating a novel approach to selectively promote vaccine induced T_SCM_ cell formation.

**Figure 7.**
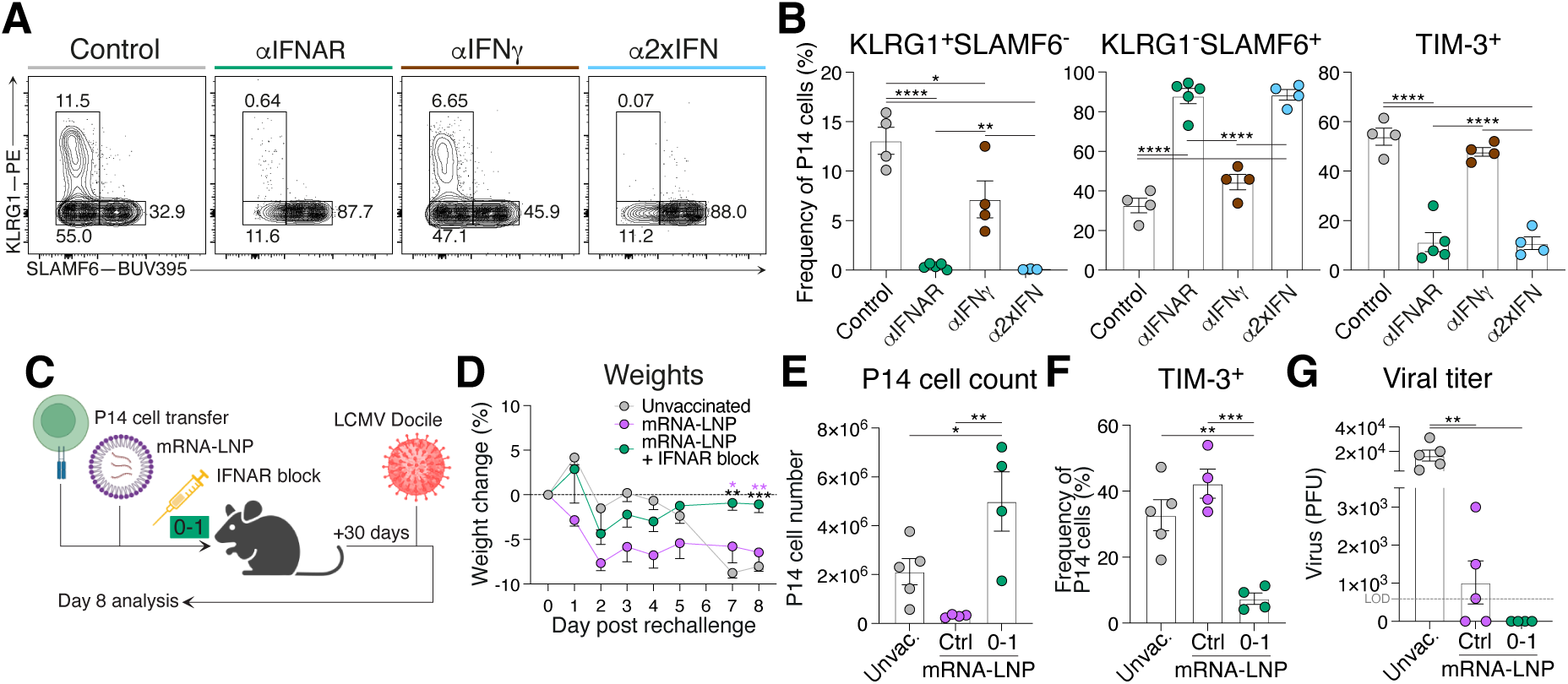
IFN blocking in combination with mRNA-LNP vaccination promotes T_SCM_ differentiation conferring enhanced protective capacity. (A, B) P14 cells from mice which received intramuscular vaccinations of GP33-encoding mRNA-LNP and d0-1 treatment of IFNAR and/or IFNγ blocking. Draining lymph node P14 cells were analyzed d8 following vaccination. Data representative of 2 independent experiments with 4-5 mice per group in each experiment. Each dot in (B) represents a single mouse. Data are mean ± S.E.M. Analyzed using one-way ANOVA tests. (A) Representative plots of P14 cells showing T_EFF_ (KLRG1^+^SLAMF6^-^), T_SCM_ (KLRG1^-^SLAMF6^+^) and T_EX_ (KLRG1^-^SLAMF6^-^TIM-3^+^) populations. (B) Graph summarizing frequencies in (A). (C-G) Comparison of response to chronic infection rechallenge in mice which received d0-1 IFNAR blocking following mRNA-LNP vaccination to unvaccinated and mice which received only vaccination. (C) Schematic illustration of experiment timeline. P14 cells were transferred into wild type hosts prior to intramuscular vaccination of GP33-encoding mRNA-LNP. Half of the vaccinated mice immediately received IFNAR, blocking followed by a secondary treatment a day later. Mice were left for 30 days to establish memory. Another cohort of naïve mice received adoptive cell transfer of naïve P14 cells a day prior to all mice being infected with chronic LCMV Docile. Mice were weighed daily following rechallenge and lymph nodes were collected d8. (D) Graph summarizes average proportion of weight loss or gain over the course of chronic LCMV infection within each group. (E) Graph of total P14 cell count in each indicated experimental group. (F) Expression of TIM-3 on the surface of P14 cells. (G) Graph summarizing the plaque forming units (PFU) from viable virus in spleens of mice in each infection condition. LOD dashed line indicates viral plaque limit of detection. *p<0.05, **p<0.01, ***p<0.001, ****p<0.0001 Supplementary Figure 6 and Supplementary Figure 7 show additional supporting data.

### IFNAR inhibition with mRNA-LNP vaccination confers superior immune protection

The early generation of T_SCM_ cells with mRNA-LNP vaccination correlates with durability of CD8^+^ T cell responses^6,34,75^. Therefore, we next sought to determine whether the near exclusive generation of T_SCM_ cells in d0-1 IFNAR blocked mRNA-LNP vaccinated mice provided improved immune recall. For this GP33 encoded mRNA-LNP vaccinated mice were rested for 30 days to establish memory. An additional cohort of unvaccinated mice received cell transfers of naïve P14 cells -1d prior to all groups being challenged with chronic LCMV Docile (Figure 7C). Consistent with chronic infection, all groups exhibited weight loss by d4-5 (Figure 7D). While the weights of unvaccinated and control vaccinated mice continued to decrease, mice that received d0-1 IFNAR blockade at the time of mRNA-LNP vaccination rapidly returned to their pre-challenge weight (Figure 7D). Despite the frequencies of T_EFF_ and T_SCM_ cell populations being similar between groups, improved recovery in d0-1 IFNAR blocked mice was associated with increased lymph node P14 cell number suggesting increased proliferative burst or increased cell survival (Figures 7E and S7A-F). Consistent with increased protection, the d0-1 IFNAR group P14 cells had reduced TIM-3 expression (Figures 7F and S7G-H) and more rapid viral clearance (Figure 7G). Therefore, we demonstrate that enhanced T_SCM_ cell differentiation leads to enhanced vaccine protection during subsequent challenges. Collectively, these data demonstrate the application of early IFNAR inhibition in a vaccine setting to drive the selective generation of T_SCM_ cells for enhance immune memory.

## DISCUSSION

The promotion of TCF-1^+^ CD8^+^ stem-like T cells is a major goal of prophylactic vaccines that elicit T cell memory and cancer therapeutic vaccines that promote tumor clearance^1,21,22^. Here, we propose that early IFN-I blockade at the time of T cell priming optimizes the formation of TCF-1^+^ T_SCM_ cells *in vivo*. Given the essential roles IFN-I plays during anti-viral responses, it may seem counterintuitive to block this pathway to enhance immune protection^39,46^. Indeed, previous work has demonstrated that IFNAR deficiency or extended blocking of IFNAR establishes persistent viral load and chronic-like infection^47,48^. Our study differs from these by limiting the IFNAR blockade to the initial days of infection, allowing viral load to be cleared. We show that blocking IFNAR during this limited window, extends the time of viral clearance, however, viral load does not reach that observed in chronic infection and is cleared within 14 days. Further, our data propose that the requirement for IFN-I to drive effector T cell differentiation is not required in vaccine settings, where there is no pathogen to overcome. Thus, impaired effector cell formation appears irrelevant to the establishment and function of vaccine-induced T cell memory.

Herein, we clarify the distinction between T_SCM_ and T_PEX_ stem-like cellular states, inferring that the presence of antigen load discriminates these two states, and that once cleared, T_PEX_ cells transition into T_SCM_ cells to maintain the memory pool. These observations are directly applicable to cancer biology and autoimmunity. Our discovery of transcriptional and cell surface molecules that distinguish these will enable the tracking of these individual cellular states during vaccination and immunotherapy^76,77^. Accordingly, we validated that CD61 and CD55 distinguish T_PEX_ and T_SCM_ cells, respectively. Interestingly, the TCF-1^+^SLAMF6^+^ CXCR6^-^CD62L^+^CD61^+^CD55^-^ T_PEX_ cell population was observed at d8 in acute LCMV Armstrong infection, suggesting that the transition from T_PEX_ to T_SCM_ cell states may naturally occur in acute infection and vaccination settings. This analysis confirmed that early IFNAR blockade results in a TCF-1^+^SLAMF6^+^CXCR6^-^CD62L^+^CD61^-^CD55^+^ T_SCM_ cell state following viral clearance. This data suggests that the T_PEX_ cellular state may reflect a natural precursor of T_SCM_ cells and this transition is interrupted in the presence of chronic viral load. Importantly, our study is distinct from other studies that block IFN-I or CXCR3 ligands in chronic infection, as we focus on IFNAR inhibition to prevent initial IFN-I signaling, rather than blocking an already chronic environment or inducing this state^47,48,50,78^. Further, our data are consistent with studies suggesting that pre-existing IFN-I hypo-responsiveness is a predictor of long-term patient survival, and suggests this is conducive to establishing an increased pool of TCF-1^+^ T cells that readily respond to PD-I blockade^79^.

Our results reveal an underappreciated interplay between type I inhibition and increased type II production, that was identified due investigation of chemokine biology. During viral infection and autoimmune disease, IFN-I and II induced signatures overlap and are thus difficult to dissect^39,51,57^. We reveal that IFN-I inhibition leads to an unexpected increase in IFN-signature chemokines. Here, we have focused on the expression of CXCL9 and CXCL10, as C57BL/6 mice lack a functional CXCL11 protein, and when CXCL11 is present there is no overt impact on CD8^+^ T cell differentiation in LCMV infection^80^. Increased *Cxcl9* and *Cxcl10* is in line with observations during infection and vaccination where IFN-I signaling is suppressed and chemokine transcription remained elevated^60,81,82^. We reveal that this increase in chemokine expression is due to IFNγ, as double deficient 2xIFN^-/-^ mice failed to induce *Cxcl9* and *Cxcl10*. The increase in IFNγ is likely due to an increase in the number of IFNγ^+^ NK cells, which may act to generate a feed-forward inflammatory loop to increase *Cxcl9* expression^55^. These results propose a re-evaluation of the influence of IFNγ in studies investigating roles of IFN-I, via IFNAR blocking or deficiency, in vaccine and viral responses and has implications for individuals with inborn errors in IFN-I signaling or neutralizing IFN-I autoantibodies^40–42,44^.

We showed that the promotion of T_SCM_ by IFNAR inhibition, was indirect of CD8^+^ T cell signaling. This is in contrast to previous studies that explored the role of IFNAR directly on CD8^+^ T cells for survival and effector formation^39,46,83,84^. We demonstrate that the promotion of stem-like T cells can be associated with increased viral load, however this is not always the case, such as with heat inactivated virus, mRNA-LNP vaccination and in *Ifng*^-/-^ hosts. Therefore, an important aspect of our study was to investigate the lymph node microenvironment and mechanism by which chemokine gradients direct T cell position cell. We demonstrate T_SCM_ cell differentiation is promoted by both low expression and overexpression of chemokines. In each of these conditions, chemokine gradient was disrupted, leading to an increase of CD8^+^ T cells in the lymph node paracortex. Although it remains unclear why paracortex positioning benefits stem-like T cell differentiation, to our knowledge, this is the first *in vivo* demonstration where inflammation leads to the abundance of chemokine which disrupts chemokine gradient formation and restricts cell migration. Our observations are consistent with the analysis of *in vitro* migration assays where desensitization of receptors limits migration potential^70^. This phenomenon of *in vivo* chemokine abundance regulating cell position may be relevant to other settings including lymphocyte congestion that is observed in the deficiency of atypical chemokine receptors or over expression of chemokines within the tumor microenvironment^85,86^.

Our results have significant implications for vaccination and adjuvant approaches that induce IFN-I and IFN-II, suggesting that in order to promote CD8^+^ memory formation, this induction should be limited^60,73,74,87,88^. We note that early IFN-I blockade decouples T_SCM_ differentiation and cell expansion, similar to other studies where stem-like T cells are promoted^89^. This is a particularly beneficial outcome in the context of vaccination to further increase the pool of memory T cells. We demonstrate that early IFNAR inhibition in combination with mRNA-LNP vaccination promotes T_SCM_ cell formation and provides a protective benefit against chronic LCMV infection, compared to mRNA-LNP vaccination alone. Further, early blocking of IFN-I signaling resembles the delayed IFN-I response observed following the YF-17D yellow fever vaccine which induces a robust and stable T_SCM_ cell population and decades-long protection^3–6^. Our data suggest the decreased IFN-I signaling early following YF-17D vaccination plays a causative role towards the preferential T_SCM_ cell induction and durability in this vaccine. Supporting this notion, vaccine responses in individuals with inborn errors in IFN-I signaling or neutralizing IFN-I autoantibodies demonstrate this is unlikely to come at a cost to humoral immunity^90^. The ability to pharmacologically induce TCF-1^+^ CD8^+^ T cells has implications for the design of vaccines to establish long-term prophylactic protection and for personalized cancer vaccines with the potential to overcome insensitivity to PD-1 blockade^1,17,21,22,91,92^. Combined, this study contributes to the basic understanding of the processes leading to T cell memory and applies this knowledge to reveal a promising approach to increase immune protection following vaccination.

## SUPPLEMENTARY FIGURES

**Supplementary Figure 1.**
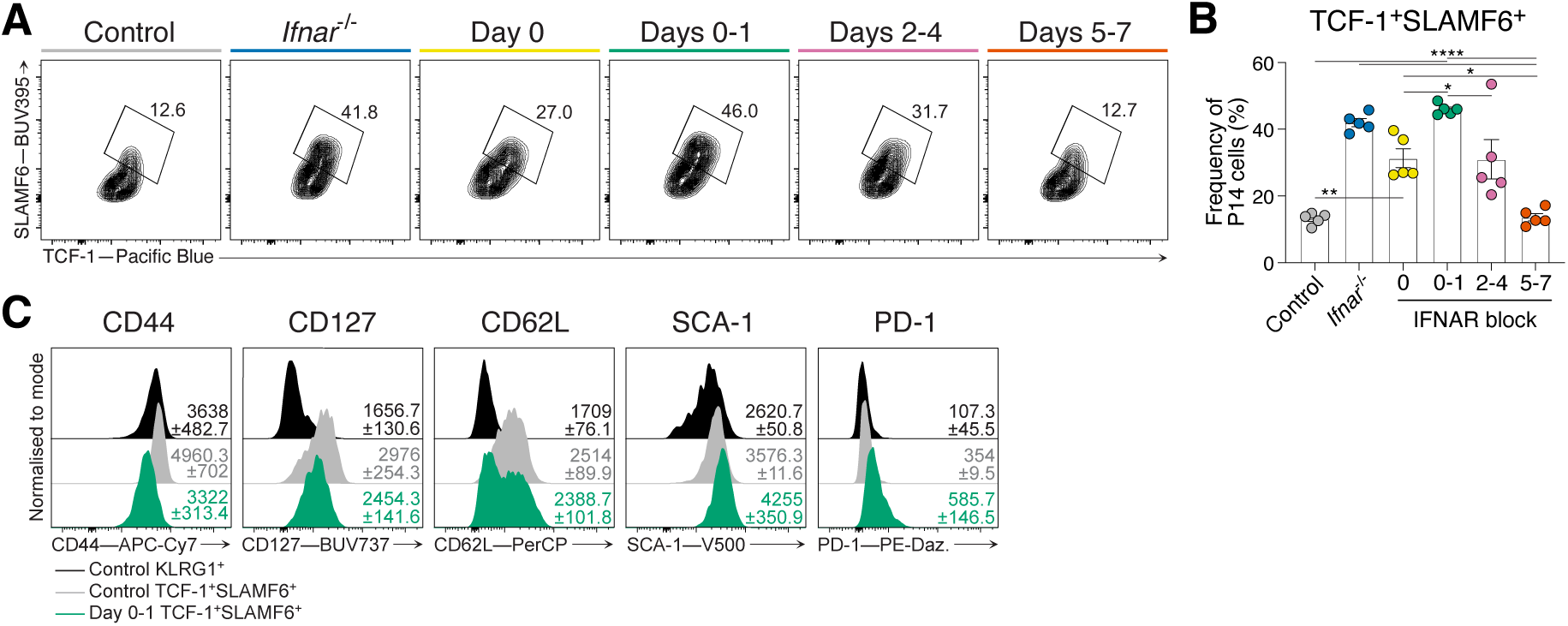
IFNAR blocking at d0-1 of acute LCMV Armstrong infection directs stem-like T cell differentiation, independent of intrinsic P14 cell IFNAR signaling. Related to Figure 1. P14 cells generated in groups indicated in Figure 1 (A). Data representative of 2 independent experiments with 5 mice per group in each experiment. Each dot in (B) represent a single mouse. Data are mean ± S.E.M. Analyzed using one-way ANOVA tests. (A) Representative plots of P14 cells showing stem-like (TCF-1^+^SLAMF6^+^) T cell populations. (B) Graph summarizing frequencies in (A). (C) Representative histograms of T_EFF_ (KLRG1^+^; black histograms) and stem-like (TCF-1^+^SLAMF6^+^; grey histograms) P14 cell populations from treated control mice and stem-like (TCF-1^+^SLAMF6^+^; green histograms) P14 cells from d0-1 IFNAR blocked mice for expression of CD44, CD127, CD62L, SCA-1 and PD-1. Data representative of 3 independent experiments with 4 mice per group in each experiment. Average gMFI ± S.E.M for each graph is indicated. *p<0.05, **p<0.01, ***p<0.001, ****p<0.0001

**Supplementary Figure 2.**
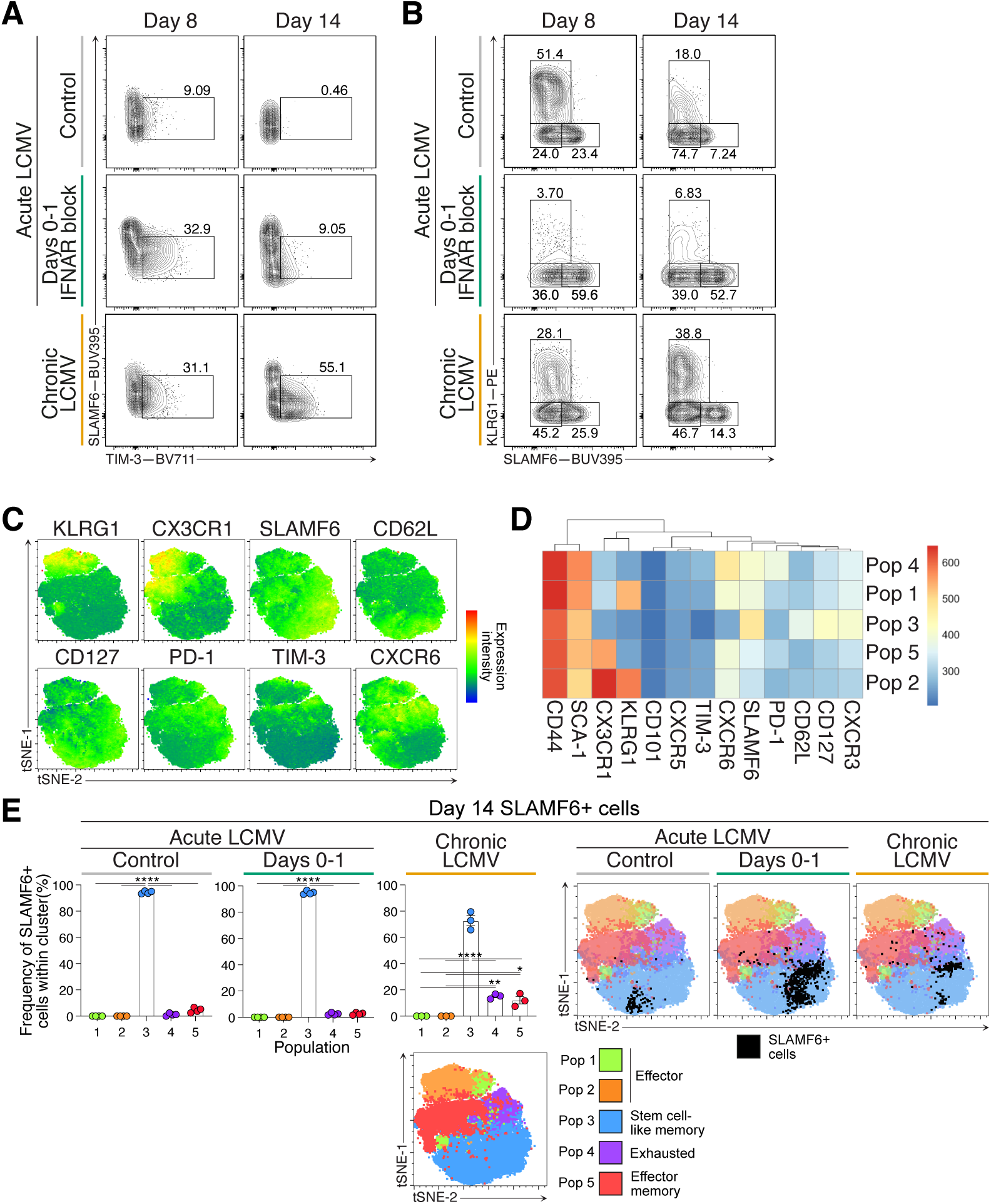
Early IFNAR blocking skews T_SCM_ cell differentiation without establishing chronic infection and exhaustion. Related to Figure 2. Analysis of P14 cells from peripheral lymph nodes of mice at d8 or d14 of acute LCMV Armstrong with or without IFNAR block at d0-1, or chronic LCMV Docile infection. Data representative of 3 independent experiments with 4 mice per group in each experiment. Each dot in (E) represent a single mouse. Data are mean ± S.E.M. Analyzed using one-way ANOVA tests. (A) Representative plots of TIM-3 expression on P14 cells within each infection condition. (B) Representative flow cytometry plots of T_EFF_ (KLRG1^+^SLAMF6^-^) and stem-like (KLRG1^-^ SLAMF6^+^) T cell populations within P14 cells from each group. (C) Overlay of marker expression heat maps on tSNE plot generated by FlowSOM. (D) FlowSOM heat map determining distinction of discreet populations. (E) Frequency of each FlowSOM population within d14 SLAMF6^+^ P14 cells for each infection condition, and corresponding representative overlay of SLAMF6^+^ P14 cells displayed in tSNE plots. *p<0.05, **p<0.01, ****p<0.0001

**Supplementary Figure 3.**
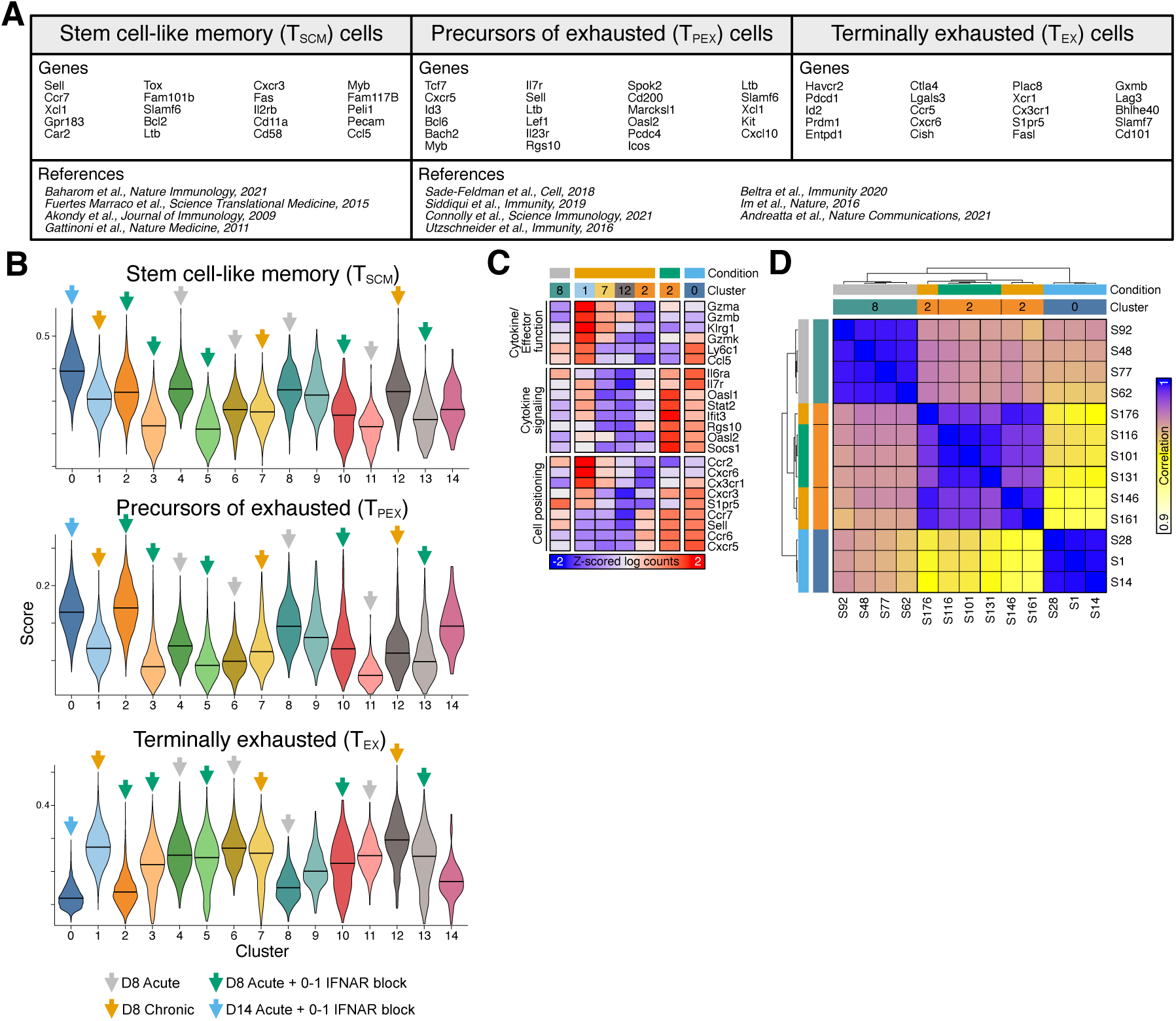
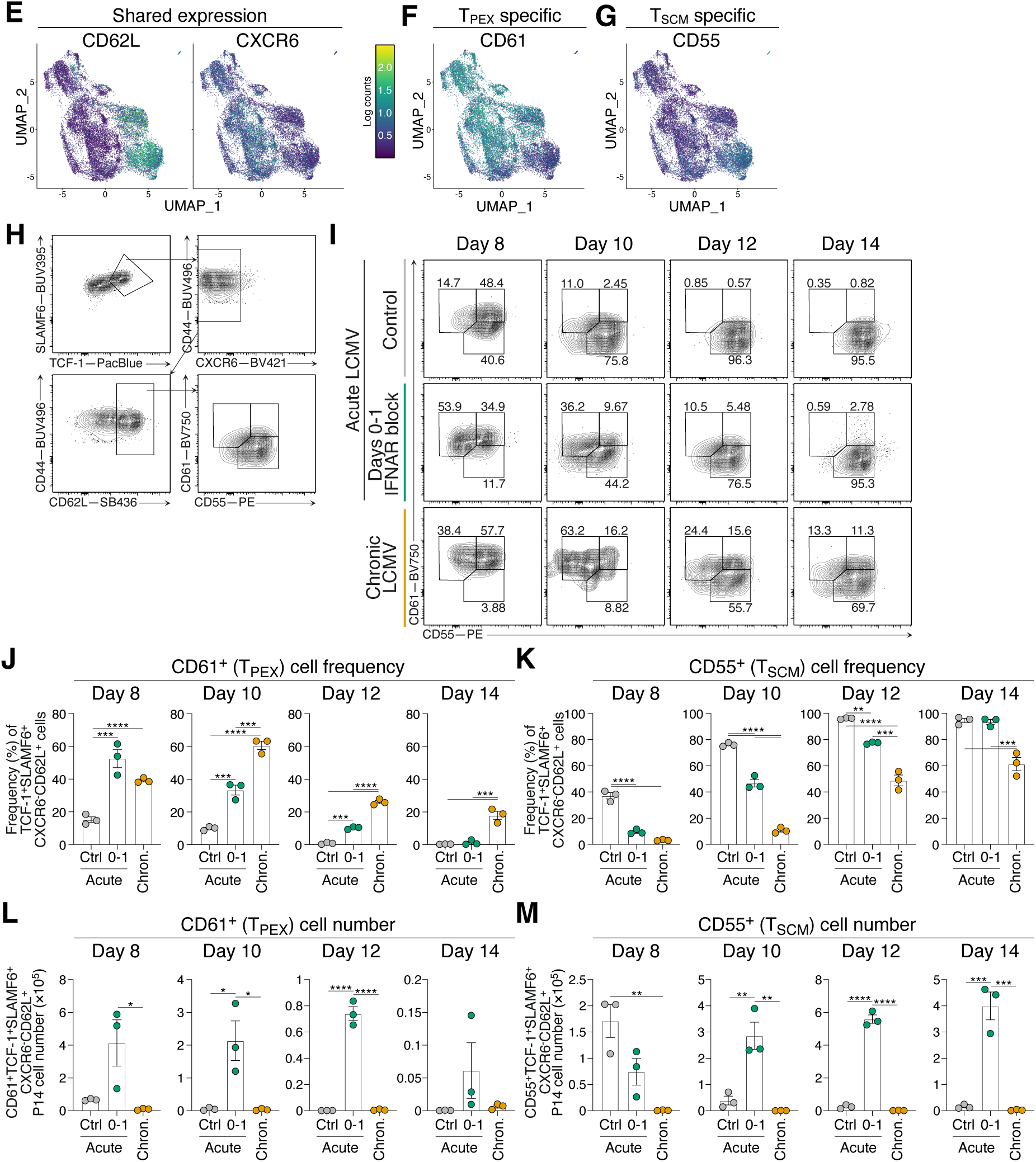
CD8^+^ T cells transition through T_PEX_ state before forming T_SCM_ cell population. Related to Figure 3. (A-G) scRNAseq and CITEseq analysis of 17629 P14 cells from peripheral lymph nodes of d8 acute LCMV Armstrong, d8 chronic LCMV Docile, d8 and d14 IFNAR blocked acute LCMV Armstrong infection conditions. Data shows 3-4 biological replicates per condition. (A) Anchor genes and reference list used to define T_SCM_, T_PEX_ and T_EX_ cell states. (B) Module scores of T_SCM_, T_PEX_ and T_EX_ cell genes within each defined cluster (from Figure 3 (A)). Prominent clusters in each experimental setting indicated with arrows. Clusters without arrows indicates clusters comprised of mixed conditions (as in Figure 2 (B)). (C) Normalized mean expression heatmap of classed marker in selected clusters. (D) Pearson’s correlation analysis of all genes expressed in pseudo-bulk samples from indicated clusters and conditions. (E-G) UMAP projections of CD8^+^ T cells showing enrichment of CITEseq defined surface marker expression. (E) Surface expression of CD62L and CXCR6 on UMAP projections of CD8^+^ T cells. (F) Surface expression of CD55 on UMAP projections of CD8^+^ T cells. (G) Surface expression of CD61 on UMAP projections of CD8^+^ T cells. (H-M) P14 cell analysis at d8 to d14 of acute LCMV Armstrong, with or without IFNAR blocking at d0-1, or chronic LCMV Docile infection. Data representative of 2 independent experiments with 6 mice per infection setting per time point. Each dot in (J, K) represent a single mouse. Data are mean ± S.E.M. Analyzed using one-way ANOVA test. (H) Gating strategy to identify TCF-1^+^SLAMF6^+^CXCR6^-^CD62L^+^CD61^+^CD55^-^ T_PEX_ cells and TCF- 1^+^SLAMF6^+^CXCR6^-^CD62L^+^CD61^-^CD55^+^ T_SCM_ cells. Pre-gated on P14 cells. (I) Representative flow cytometry plots to identify frequencies of TCF-1^+^SLAMF6^+^CXCR6^-^ CD62L^+^CD61^+^CD55^-^ T_PEX_ cells and TCF-1^+^SLAMF6^+^CXCR6^-^CD62L^+^CD61^-^CD55^+^ T_SCM_ cells. (J, K) Graphs summarizing (I) throughout infection. (L, M) Graphs summarizing the cell numbers of (I) throughout infection. **p<0.01, ***p<0.001, ****p<0.0001

**Supplementary Figure 4.**
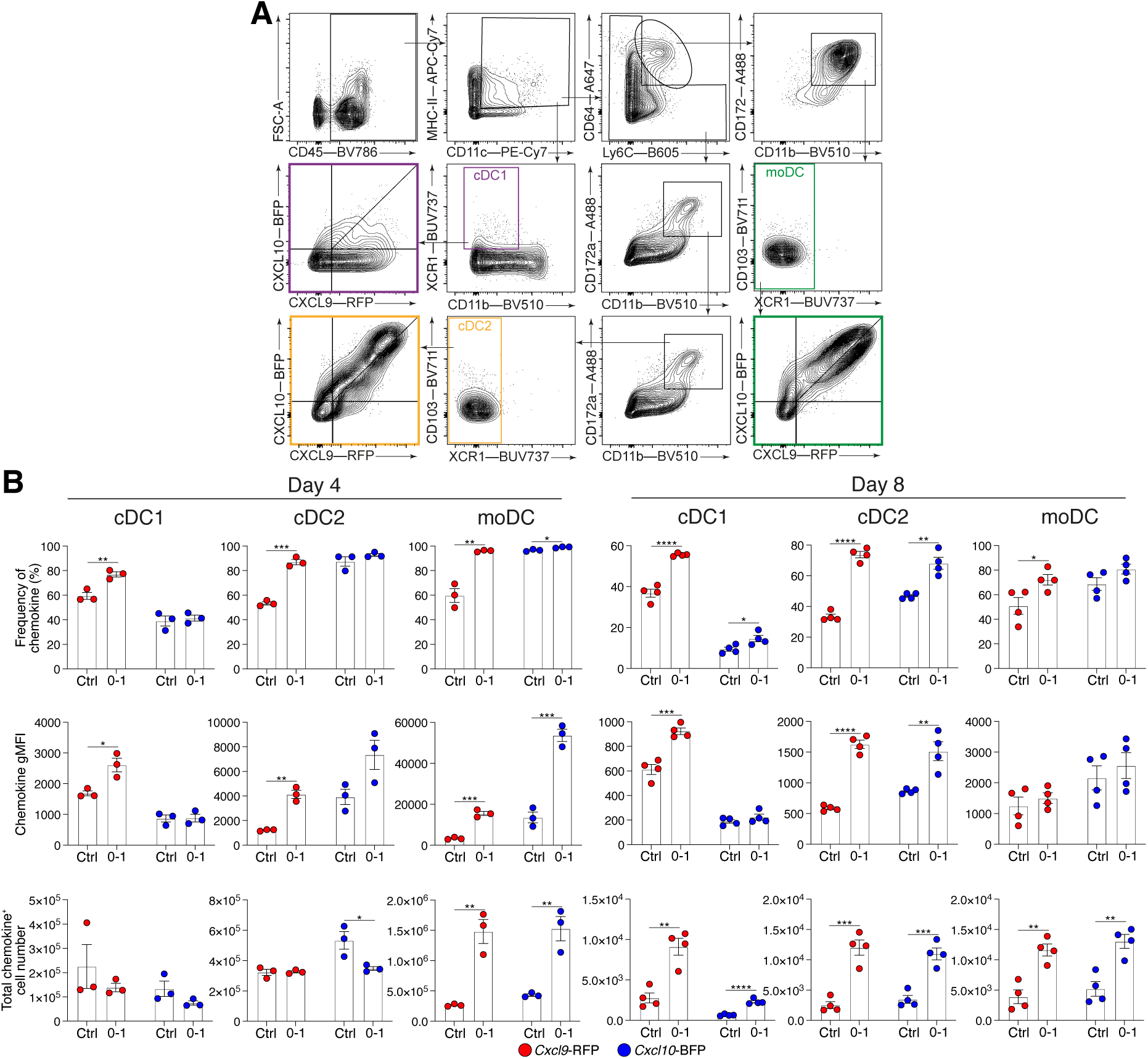
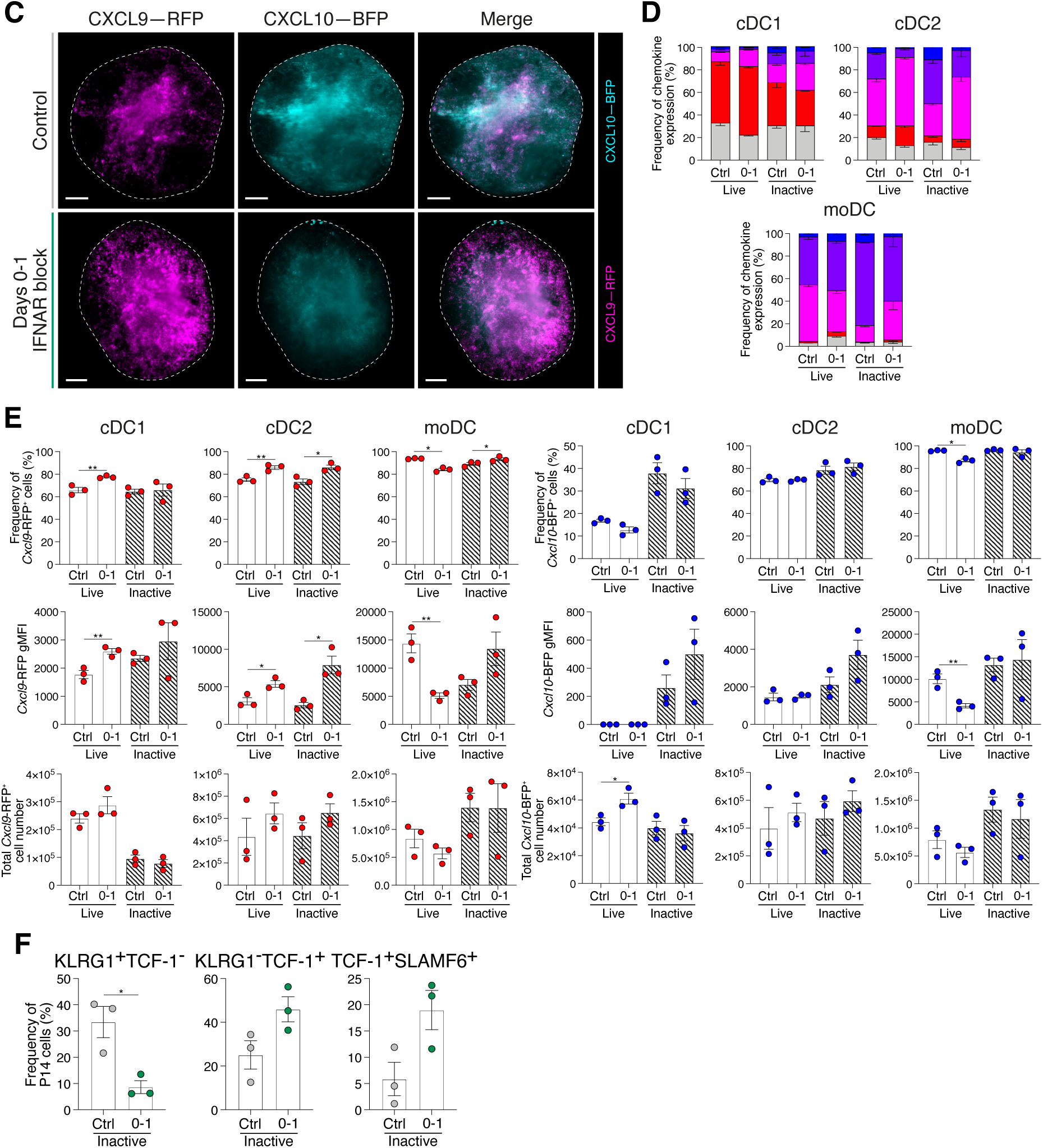
DC gating strategy and increased CXCR3 ligand expression and T_SCM_ cell differentiation during live and heat inactivated acute LCMV Armstrong infection following IFNAR blocking. Related to Figure 4. (A) Gating strategy to identify cDC1, cDC2 and moDC cell populations. Colored boxes indicate final gate and REX3 expression for each subset. (B) Frequency, gMFI and total cell number of *Cxcl9*-RFP^+^ and *Cxcl10*-BFP^+^ DC subsets at d4 and d8 post LCMV Armstrong infection. Data are representative of 3 individual experiments with 3-4 mice in each group. (C) LSFM micrographs of intact REX3 lymph nodes d8 of infection in control and d0-1 IFNAR blocked conditions. Images are 200µm longitudinal slices through lymph node center. Scale bars represent 100µm. Dashed line indicates lymph node outline. Individual REX3 reporters and merge images show *Cxcl9*-RFP (magenta), *Cxc10*-BFP (cyan). Images are representative of 3 individual experiments with at least 4 mice in each group in each experiment. (D, E) Analysis of *Cxcl9*-RFP and *Cxcl10*-BFP expression in DC subsets within peripheral lymph nodes of mice d8 following challenge with either live or heat inactivated (inactive) LCMV Armstrong. Half of each group received d0-1 treatments of IFNAR block. Data representative of 3 individual experiments with 3 mice per group per experiment. Data are mean ± S.E.M. (D) Graphs summarizing the graded frequencies *Cxcl9*-RFP^+^ and *Cxcl10*-BFP^+^ DC subsets d8 following acute LCMV Armstrong infection (E) Frequency, gMFI and total cell numbers gMFI of chemokine reporter expression in each immune challenge condition. Mean ± S.E.M. Analyzed using unpaired t-tests. Each dot in (B, E, F) represents a single mouse. Mean ± S.E.M. Analyzed using unpaired t-tests. *p<0.05, **p<0.01, ***p<0.001, ****p<0.0001

**Supplementary Figure 5.**
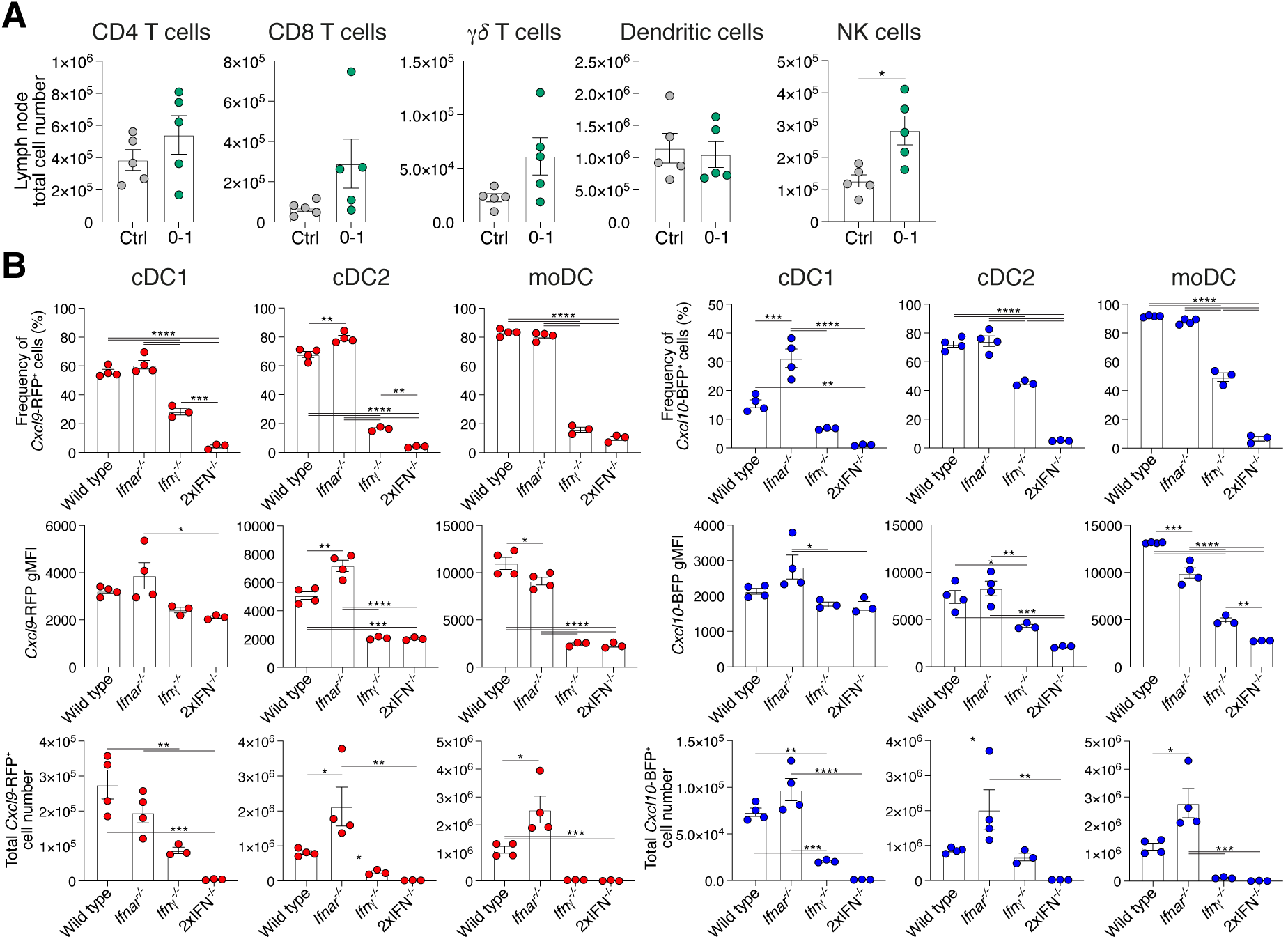
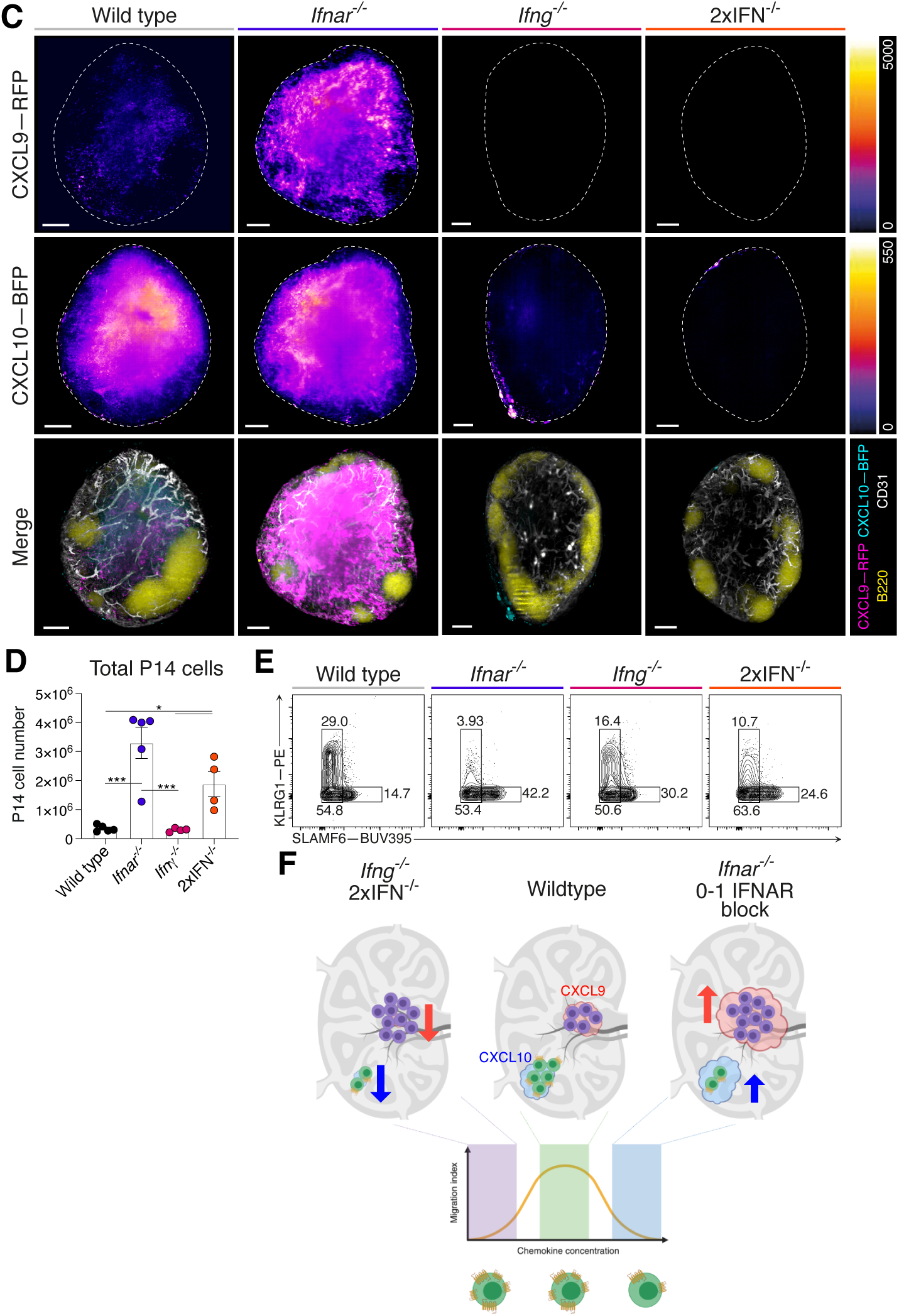
Absence of IFN-I and IFN-II signaling promotes T_SCM_ cell differentiation during acute LCMV Armstrong infection. Related to Figure 6. (A) Total lymph node IFNg^+^ of indicated cells from control treated and d0-1 IFNAR blocked mice at d4 of acute LCMV Armstrong infection. (B) Expression of *Cxcl9*-RFP and *Cxcl10*-BFP in cDC1, cDC2 and moDC populations from REX3 hosts crossed to indicated IFN deficient mice. (C) LSFM micrographs of intact wild type, *Ifnar^-/-^*, *Ifng^-/-^*, and 2xIFN^-/-^ mice REX3 lymph nodes. Images are 200µm longitudinal slices through lymph node center. Scale bars represent 100µm. Dashed line indicates lymph node outline. Individual REX3 reporters and merge images show *Cxcl9*-RFP (magenta), *Cxcl10*-BFP (cyan). Images are representative of 3 individual experiments with at least 3 mice in each group in each experiment. (D) Total P14 cell number in each wild type of IFN deficient host mouse. (E) Representative plots of P14 cells showing T_EFF_ (KLRG1^+^SLAMF6^-^), T_SCM_ (KLRG1^-^SLAMF6^+^) and T_EX_ (KLRG1^-^SLAMF6^-^) cells. (F) Model of IFN control of chemokine production and CD8^+^ T cell position within lymph nodes and how this correlates with *in vitro* migration assay chemokine concentration. Lymph nodes indicate P14 cell location and chemokine receptor expression in *Ifng^-/-^* and 2xIFN^-/-^ (left), wildtype (middle), and *Ifnar^-/-^* and d0-1 IFNAR blocked (right) settings. Dotted lines indicate correlation to chemokine concentration and bell curve of cell migration index in *in vitro* assays. The wildtype setting correlates with optimal CXCL9 (red) and CXCL10 (blue) gradient formation to facilitate the generation of both T_EFF_ (green) and T_SCM_ (purple) cells. *Ifng^-/-^*and 2xIFN-/- settings exhibit low CXCL9 and CXCL10 expression to promote cell retention in paracortex, increasing the generation of T_SCM_ cells and reduced T_EFF_ cells. In *Ifnar^-/-^* and d0-1 IFNAR blocked settings, increased CXCL9 and CXCL10 expression causes down regulation of surface CXCR3, preventing cell migration and increased T_SCM_ cell and reduced T_EFF_ cell differentiation. Each dot in (A, B, D) represents a single sample. *p<0.05, **p<0.01, ***p<0.001, ****p<0.0001

**Supplementary Figure 6.**
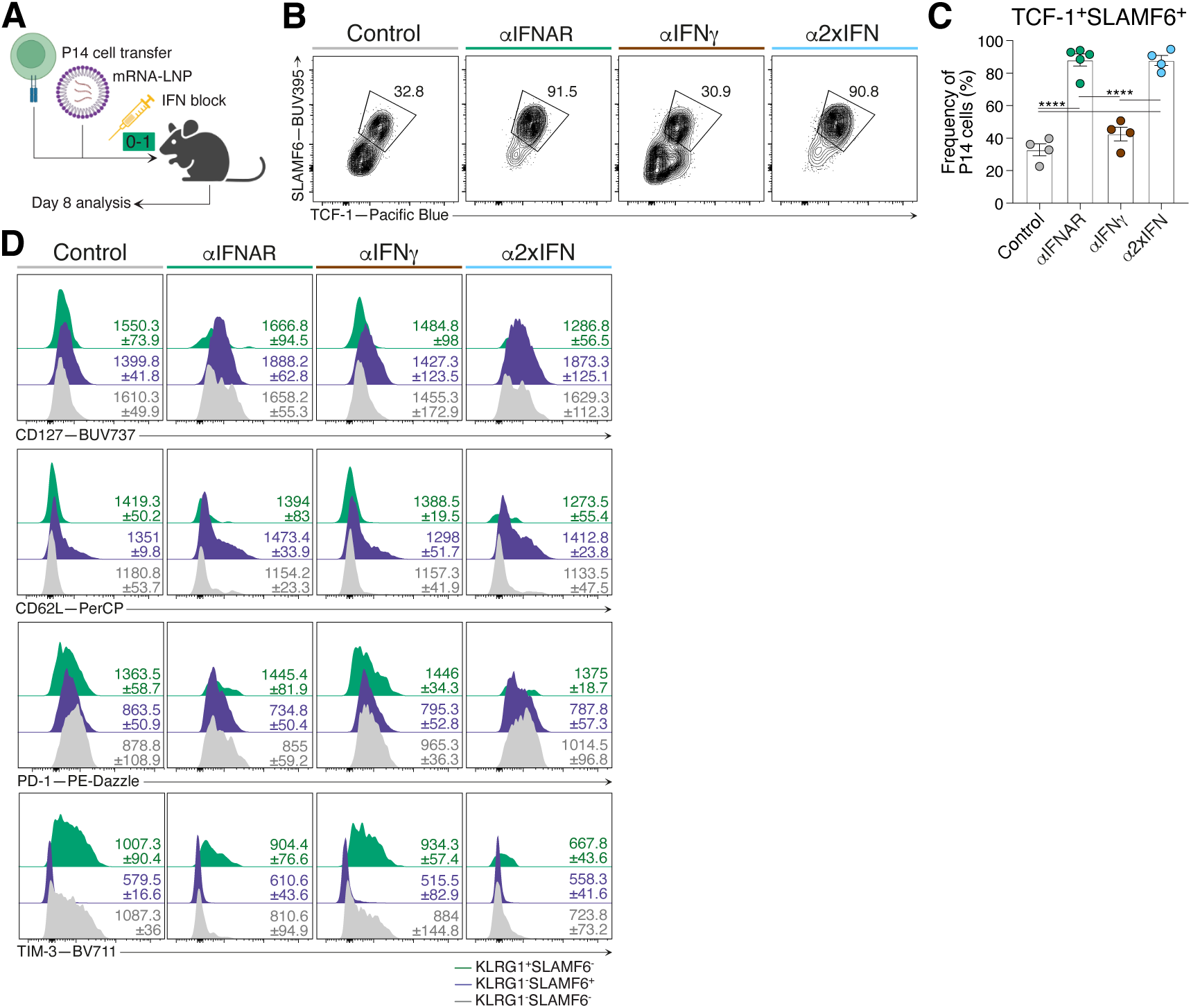
IFN inhibition alongside mRNA-LNP vaccination promotes exclusive generation of T_SCM_ cells. Related to Figure 7. Draining lymph node P14 cells from mice d8 following GP33-encoding mRNA-LNP vaccination in combination with IFNAR and/or IFNg blocking at d0-1. (A) Experimental scheme. Naïve P14 cells were adoptively transferred into wild type host mice 24 hours prior to mRNA-LNP vaccines which encode the GP33 epitope. Cohorts did not receive further treatment or received treatments of either IFNAR blocking, IFNg blocking, or a combination at d0-1 following vaccination. (B) Representative flow cytometry plots showing T_SCM_ (TCF-1^+^SLAMF6^+^) cell populations of P14 cells within each condition. (C) Graph summarizing frequencies in (B). Each dot represents a single mouse sample. (D) Representative histograms of T_EFF_ (KLRG1^+^SLAMF6^-^), T_SCM_ (KLRG1^-^SLAMF6^+^) and T_EX_ (KLRG1^-^SLAMF6^-^) P14 cell populations from control, IFNAR blocked, IFNg blocked or combined IFNAR and IFNg blocked mice for expression of CD127, CD62L, PD-1 and TIM-3. Average gMFI ± S.E.M for each graph is indicated. ****p<0.0001

**Supplementary Figure 7.**
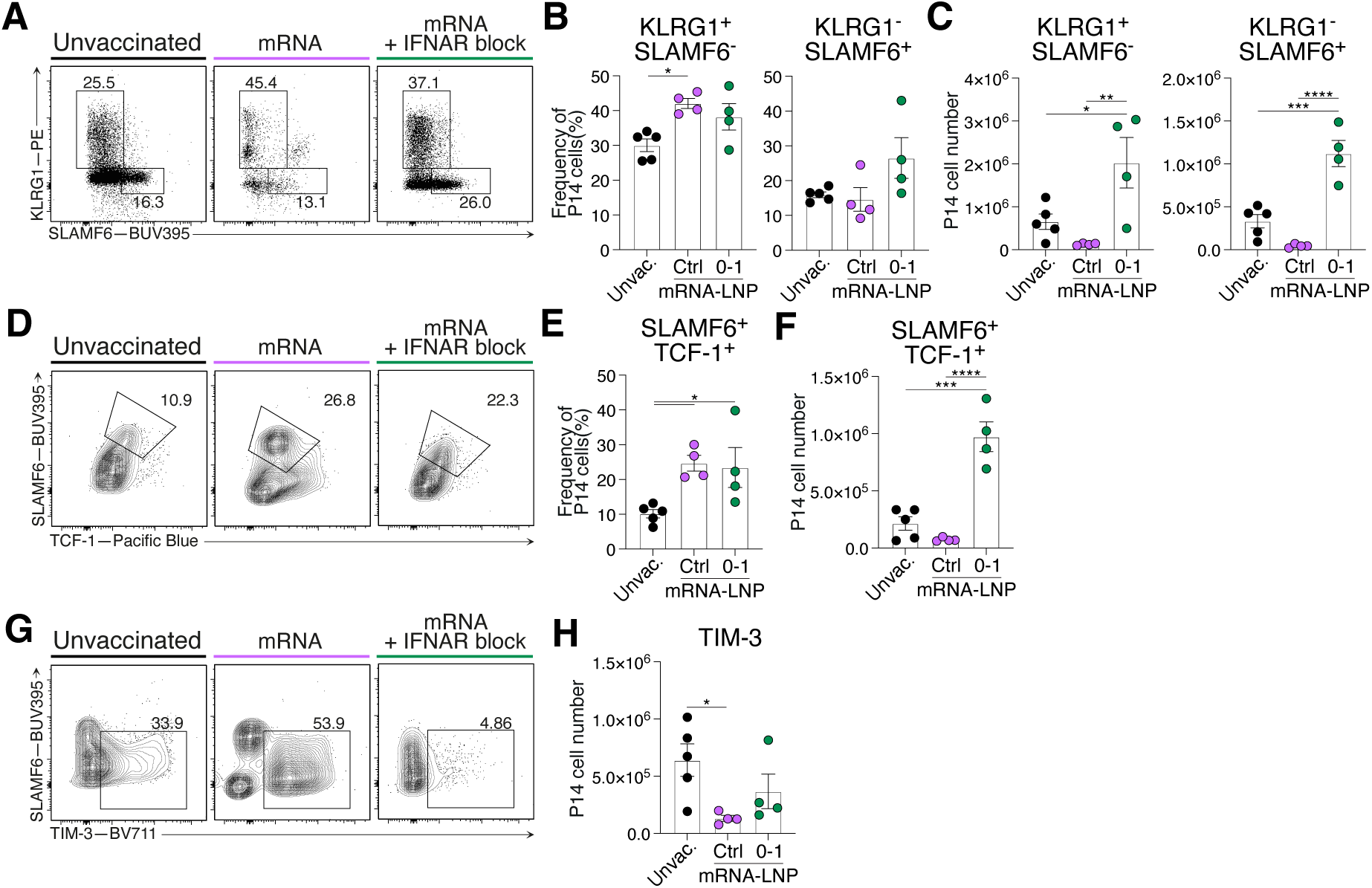
T_SCM_ cells generated following d0-1 IFNAR inhibition in combination with mRNA-LNP drive increased proliferation of specific T cells upon viral rechallenge. Related to Figure 7. P14 cell analysis d8 after rechallenge in mice which received d0-1 IFNAR blocking following mRNA-LNP vaccination to unvaccinated and mice which received vaccination. Data representative of 2 individual experiments with 4-5 mice per group per experiment. Data are mean ± S.E.M. Each dot in (B, C, E, F, H) represents a single sample. (A) Representative plots of P14 cells showing T_EFF_ (KLRG1^+^SLAMF6^-^) and T_SCM_ (KLRG1^-^SLAMF6^+^) cell populations. (B) Graphs summarizing population frequencies in (A). (C) Cell counts of showing T_EFF_ (KLRG1^+^SLAMF6^-^) and T_SCM_ (KLRG1^-^SLAMF6^+^) cell populations in each experimental condition. (D) Representative flow cytometry plots of T_SCM_ (TCF-1^+^SLAMF6^+^) P14 cell populations. (E) Graph summarizing frequencies in (D). (F) T_SCM_ (TCF-1^+^SLAMF6^+^) cell counts. (G) Representative flow cytometry plots of TIM-3 expression of P14 cells within each condition. (H) Graph summarizing frequencies in (G). *p<0.05, **p<0.01, ***p<0.001, ****p<0.0001

## RESOURCE ABAILABILITY

### Lead contact

Further information and requests for resources and reagents should be directed to and will be fulfilled by the lead contact, Joanna Groom (; @groomlab)

### Materials availability

All experimental models and reagents will be made available upon installment of a material transfer agreement.

## EXPERIMENTAL MODEL AND SUBJECT DETAILS

### Mice

Mice were bread and maintained on a C57BL/6 background under specific pathogen-free conditions. REX3 transgenic^69^, *Ifnar*^-/93^, *Ifng*^-/-94^, *Cxcl9*^-/-95^, *Cxcl10*^-/-96^, P14 transgenic^97^ and Ly5.1^98^ mice have been previously described. REX3 mice were bred with *Ifnar*^-/-^ and/or *Ifng*^-/-^ to generate REX3 *Ifnar*^-/-^, REX3 *Ifng*^-/-^, and REX3. *Ifnar*^-/-^*Ifng*^-/-^ mice. Mice with Green Fluorescence Protein (GFP) under the chicken beta-actin promoter^99^ were bred with P14 and Ly5.1 mice. All experiments were conducted in compliance with the Walter and Eliza Hall Institute Animal Ethics Committee and performed on mice of 6-10 weeks of age of mixed sex.

## METHOD DETAILS

### Adoptive cell transfer, viral infection and IFN inhibition

Naïve P14 cells were isolated using the Naïve CD8a^+^ T cell Isolation Kit (Miltenyi Biotec) and 2×10^4^ cells were transferred into host mice via intravenous lateral tail injection one day prior to viral infection. Mice were inoculated with 3×10^4^ or 2×10^6^ plaque-forming units (PFU) of LCMV Armstrong or LCMV Docile, respectively, via intravenous lateral tail injection. Unless indicated otherwise, mice received two (d0-1) intraperitoneal injections of, either or combined, 200μg anti-IFNAR monoclonal antibodies (clone MAR1-5; Leinco Technologies and gift from Paul J. Herzog) and 200μg anti IFNγ monoclonal antibodies (WEHI Antibody Facility).

### Preparation of samples for flow cytometry

Blood samples were twice treated with red cell lysis buffer (WEHI Media) for 3 minutes at room temperature to remove red blood cells. Single-cell suspensions were stained for surface antigen expression using indicated antibodies (Key Resources Table) for 20 minutes at 4°C, followed by viability dye staining for 10 minutes at 4°C. Transcription factor and chemokine staining were performed using the Foxp3 Transcription Factor Staining Kit (ThermoFisher Scientific). FlowSOM analysis defined cell clusters based on differential staining of CXCR3, CD127, CD62L, PD-1, SLAMF6, CXCR6, TIM-3, CXCR5, CD101, KLRG1, CX3CR1, SCA-1 and CD44. All flow cytometry analysis was performed on a BD LSRFortessa X-20 and BD FACSymphony A3 Cell Analyzers (BD Biosciences) data was analyzed using FlowJo v10 (FlowJo LLC) and the FlowSOM plugin.

### Preparation of samples for cell sorting and single cell RNA sequencing

Lymph nodes were harvested and processed to form a single cell suspension. Samples were stained for surface antigen expression using indicated antibodies (Key Resources Table) and TotalSeq HashTags (BioLegend) for 30 minutes at 4°C, followed by viability dye staining for 10 minutes at 4°C. Sample suspensions were sorted using BD FACSAria Fusion Flow Cytometer (BD Biosciences) to isolate P14 CD8^+^ T cells.

### Viral titre of tissues

Spleen tissues were harvested and homogenized using Qiagen TissueLyser for 7 minutes at 30Hz. LCMV viral titers were determined using a focus-forming assay, as previously outlined^100^.

### Data processing and demultiplexing of single cell RNA sequencing data

For data processing, reads from each capture were processed using 10X Genomics Cell Ranger software (v7.0.0). Firstly, ‘cellranger mkfastq’ and bcl2fastq (v2.19.1) were used to convert and demultiplex the Illumina sequencer’s BCL files into FASTQ files for each of the gene expression (GEX), antibody derived tag (ADT), and hashtag oligo (HTO) libraries. Secondly, ‘cellranger multi’ was used with default settings to generate count matrices. The GEX data were mapped and quantified against the 10X Genomics pre-built mm10 (GENCODE vM23/Ensembl 98) reference groups and transcriptome (2020- A (July 7, 2020) version) and the feature barcoding (ADT and HTO) data were quantified against a ‘feature.csv’ file containing the barcode sequences provided by BioLegend. Finally, the DropletUtils R/Bioconductor package (v1.18.1) was then used to load the Cell Ranger output files into R (v4.2.1) and to identify non-empty droplets using ‘emptyDrops’ method with default settings^101^. For demultiplexing, the ‘demuxmix’ method from the demuxmix (v1.0.0) R/Bioconductor package was applied to the HTO data, with the ‘naive’ model and default parameters, to multiplex non-empty droplets to their sample of origin. This was performed separately for each capture. A droplet was assigned to a sample if the best demuxmix assignment matched the corresponding HTO combination of a sample or was otherwise assigned as a ‘multiplet,’ ‘negative,’ or ‘uncertain’ sample. All scripts used are available from https://github.com/WEHISCORE/G000304_Broomfield.

### Analysis of single cell RNA sequencing and CITE sequencing data

#### QC and clustering

QC on the demultiplex-ed scRNA-seq data was conducted using scater R package^102^ to remove low quality cells and cells with high mitochondrial genes (≥ 10%), leaving 17,629 cells for analysis. The data was then normalized using SCTransform in Seurat R package^103^ and clustered using the default Louvain method at resolution 0.8, resulting in 15 clusters. Curation based on proportions distribution by conditions led to the merging of two clusters, resulting in 14 clusters for downstream analysis. Modules scores of each cluster was calculated using the ‘AddModuleScore’ function in Seurat based on the anchor genes selected from previous studies for T_SCM_, T_PEX_, and T_EX_ cells (Supplementary Figure 3 (A)6-8,11,13,14,17,19,34,77,104.

#### Differential expression analysis

Differential expression (DE) was conducted using a pseudo-bulked approach based on “cluster” and “samples”, and after filtering out samples with less than 10 cells and applying gene level QC using edgeR::filterByExpr, leaving in 131 pseudo-samples and 9438 genes for analysis. Differential expression analyses were conducted using a *voom-limma-duplicatecorrelation with sample weights* pipeline via the *edgeR::voomLmFit* function to fit a linear model with “condition_cluster” (condition being infectious setting) as the covariate, and to estimate the consensus correlation across mice and account for mice variation as a random effect^105,106^. DE were conducted for the following comparisons based on 4 clustered identified from the cluster analysis: (A) d8 acute LCMV cluster C8 vs all other clusters, (B) d8 chronic LCMV cluster C2 vs all other clusters, (C) d8 IFNAR blocked acute LCMV cluster C2 vs all other clusters, (D) d14 IFNAR blocked acute LCMV cluster C0 vs all other clusters, and (E) clusters (d8 chronic LCMV cluster C2 + d8 IFNAR blocked acute LCMV cluster C2) vs clusters (d14 IFNAR blocked acute LCMV cluster C0 + d8 acute LCMV cluster C8). An empirical Bayes moderated t-statistic was generated with multiple testing adjustment carried out using the Benjamini–Hochberg procedure to identify statistically significant genes (adjusted *P* < 0.05). The enrichment of marker gene sets from published datasets^6,8,11,17,19,34,66,107,108^ in the DE results was calculated using the *fgsea* package^109^ and visualized as normalized enrichment score (NES) dot plots.

#### Data processing and visualization of CITE-seq analysis

CITE-seq data was extracted and normalized using the CLR (centered log ratio transformation) method across cells from the *Seurat::NormalizeData* function. The logcount expression for selected markers were visualized by overlaying onto the scRNA-seq UMAPs of CD8^+^ T cells ^110^.

### Dendritic cell isolation from lymph nodes

Lymph nodes were mechanically disrupted with tweezers and incubated in RPMI with 0.8mg/mL Dispase II (Roche), 0.2mg/mL Collagenase P (Roche), and 0.1mg/mL DNase I (Sigma-Aldrich) for 20 minutes at 37°C. Supernatant was collected, and remaining tissue pieces were further incubated twice more in fresh digestion medium for 10 minutes at 37°C each.

### Whole tissue immunofluorescence staining, clearing, light sheet fluorescence microscopy (LSFM)

Lymph nodes were harvested and fixed in 4% Paraformaldehyde (Sigma-Aldrich) for 12-24 hours at 4°C, followed by incubation in blocking buffer containing 1% Bovine serum albumin (Sigma-Aldrich), 1% Normal rat serum (Jackson ImmunoResearch) and 0.3% Triton X-100 (Sigma-Aldrich) in PBS for 24 hours at 4°C. Whole tissues were stained with indicated antibodies in blocking buffer for three days at 4°C. Tissues were washed by immersion in PBS containing 0.5% 1-thioglycerol (Sigma-Aldrich) and 0.2% Triton X-100 (Sigma-Aldrich) for 24 hours at room temperature. Lymph nodes were immersed in Ce3D clearing medium containing 1.455g/mL Histodenz (Sigma-Aldrich), 40% N-methylacetamide (Sigma-Aldrich), 0.5% 1-thioglycerol and 0.1% Triton X-100 in PBS for 24 hours at room temperature on a shaking incubator^111^. The clearing medium was replaced with fresh medium and tissues were incubated for a further two-three days to clear the lymph nodes to a refractive index of 1.49-1.5. The cleared lymph nodes were embedded in 2% low-melting agarose (Sigma-Aldrich) containing 1:10,000 Fluoresbrite YG miscrospheres 1µm (Polysciences) in 2.15mm diameter glass capillaries (Zeiss). Samples were submerged in clearing solution for 24 hours before imaging to allow for refractive index matching between the agarose and clearing medium. For light sheet fluorescence microscopy, images were acquired on a Z.1 Light sheet microscope (Zeiss) using a 5x (f/0.16) air objective. LSFM images were processed using Zen Blue and Zen Black (Zeiss) and Imaris 9.7.2 (Oxford Instruments). Pseudocolor FIRE intensity LUT scales were set to brightest pixel intensity across samples and maintained for all comparisons for each reporter protein.

### Quantification of cells in intact lymph nodes

Analyses performed using the eroded volume fraction (EVF) implemented within the 3D ImageJ Suite v2.1.0/1.53^36,112^. First, lymph node images were smoothed using a 3D median filter (with radius rx = 4, ry = 4 and rz = 2). 3D lymph nodes were then segmented using a manually set global thresholding. Images containing cell signals were filtered with a 3D median filter (with radius rx = 2, ry = 2 and rz = 1) followed by a top-hat filtering (with radius rx = 6, ry = 6 and rz = 3) to enhance the signal of spots. The cell signal was manually set using global thresholding. The positioning of cells within the lymph nodes was assessed using an EVF analysis, whereby a 3D distance map was computed inside the lymph node, then the distance values were sorted and normalized from 0 to 1. The EVF values were then divided into 100 layers of equal volumes from 0 (near the lymph node periphery) to 1 (lymph node center), and the volume of cells within each layer was computed.

### Migration assay

Axillary, brachial and inguinal lymph nodes were harvested and mashed into single cell suspension. Then, 4 × 10^6^ total cells were added to 96-well transwells (Corning Costar) in RPMI with 0.5% FCS and migrated toward 1 to 10,000 ng/ml CXCL10 chemokine (PeproTech) for 1.5 hours. Cells in the lower chamber were collected and stained. Migration of P14 cells was detected by timed acquisition on a BD FACSymphony A3 Cell Analyzer (BD Biosciences). The migration index was defined as the percentage of migrated cells relative to the input sample.

### IFNγ cytokine bead array on lymph node tissue lysates

Lymph nodes were harvested from d4 infected mice and snap frozen on dry ice. The tissue was weighed and digested with 10μL mg^−1^ DISC lysis buffer (20 mM Tris-HCl pH 7.5, 150 mM NaCl, 2 mM EDTA, 1% TritonX-100, 10% glycerol, H_2_O, protease inhibitor cocktail tablet: Roche, Basel, Switzerland). The tissue was lysed with TissueLyser85300 (Qiagen, Hilden, Germany) and the supernatant was collected between the pellet and fat layers following a 20 000 *g* 15 min spin. Supernatant from tissue lysate was diluted 1:10 and loaded onto BD Cytometric Bead Array Mouse IFNg Flex set (BD Biosciences, Franklin Lakes, NJ, USA), used as per the manufacturer’s instructions. Briefly, tissue lysates and Mouse IFNg standards were added to IFNg cytokine capture beads and PE detection reagent in 96-well plate for 2h at room temperature, protected from light. The plate was washed two times in BD CBA wash buffer, resuspended and acquired on a BD LSRFortessa X-20 cell analyzer (BD Biosciences). Data analysis and standard curve generation was performed with FCAP Array Software version 3.0 (BD Biosciences).

### mRNA-LNP vaccine production and vaccination

mRNA production was performed as described^113^. Briefly, the sequence of the P14 minigene was codon-optimized, synthesized (GenScript), and cloned into an mRNA production plasmid. The mRNA was produced from the linearized plasmid to contain 101 nucleotide-long poly(A) tail. m1Ψ- 5’triphosphate instead of UTP was used to generate modified nucleoside-containing mRNA. Capping of the in vitro transcribed mRNA was performed co-transcriptionally using the trinucleotide cap1 analog, CleanCap (TriLink). mRNA was purified by cellulose purification, as described^114^. The minigene-encoding mRNA was analyzed by agarose gel electrophoresis and were stored frozen at - 20°C. The cellulose-purified m1Ψ-containing mRNA were encapsulated in LNP using a self-assembly process as previously described wherein an ethanolic lipid mixture of ionizable cationic lipid, phosphatidylcholine, cholesterol and polyethylene glycol-lipid was rapidly mixed with an aqueous solution containing mRNA at acidic pH^115^. The LNP formulation used in this study is proprietary to Acuitas Therapeutics; the proprietary lipid and LNP composition are described in US patent US10,221,127. The mRNA-loaded particles were characterized and subsequently stored at -80°C at an RNA concentration of 1μg μl-1 . The mean hydrodynamic diameter of mRNA-LNP was ∼80 nm with a polydispersity index of 0.02-0.06 and an encapsulation efficiency of ∼95%.

For vaccination, mice received a single intramuscular caudal thigh injection of 10µg GP33 minigene (MKAVYNFATM)-encoding mRNA-LNP prior to intraperitoneal injections of, either or combined, 200µg anti-IFNAR or 200µg anti-IFNγ (WEHI Antibody Facility) monoclonal antibodies.

## QUANTIFICATION AND STATISTICAL ANALYSIS

Statistical differences between groups in datasets with one categorical variable were evaluated by unpaired t-tests (2 groups) or one-way ANOVA (more than 2 groups) corrected for multiple comparisons. Statistical differences between groups in datasets with two categorical variables were evaluated by two-way ANOVA corrected for multiple comparisons. Exact p values are given for statistical differences between p 0.05 and 0.0001. All experimental data are presented as mean ± standard error of the mean (S.E.M.) with statistical analysis performed using Prism 8 (GraphPad Software).

## ACKNOWLEDGEMENTS

Some anti-IFNAR antibody for *in vivo* use was gifted from Paul J. Hertzog. Paired scRNAseq and CITEseq experiments were technically supported by WEHI Genomics Facility. Experimental scheme figures were generated with BioRender. This work was supported by National Health and Medical Research Council (NHMRC) Ideas grant (1182649), and Cancer Council NSW grant (2019642) to J.R.G., B.J.B., R.Z.Q., and L.D. are supported by Melbourne research scholarship. B.C.D. is supported by WEHI Academic Excellence scholarship. J.R.G. is an NHMRC Leadership Investigator (2007812).

N.P. was supported by the National Institute of Allergy and Infectious Diseases (R01AI146101 and R01AI153064). M.P. is supported by an NHMRC Investigator (1175011). This work was supported by the Norman, Ann and Graeme Atkins Charitable Trust. This work was made possible through Victorian State Government Operational Infrastructure Support and Australian Government NHMRC IRIISS.

## AUTHOR CONTRIBUTIONS

Conceptualization: J.R.G.

Methodology: B.J.B., C.W.T., R.Z.Q., B.C.D., L.D., J.C., and V.C.W.

Investigation: B.J.B., C.W.T., R.Z.Q., B.C.D., C.A., L.D., J.C., L.M, and H.M.

Software and Formal Analysis: B.J.B., C.W.T., R.Z.Q., B.C.D., J.C. and V.C.W.

Visualization: B.J.B., V.C.W., C.W.T., and J.C.

Resources: M.P., K.L.R, H.M., N.P., W.J.M., M.J.D., and J.R.G.

Writing – Original draft and revisions: B.J.B., and J.R.G.

Funding Acquisition: J.R.G.

Supervision: J.R.G.

